# Physiological febrile heat stress increases cytoadhesion through increased protein trafficking of *Plasmodium falciparum* surface proteins into the red blood cell

**DOI:** 10.1101/2025.05.14.653932

**Authors:** David Jones, Hugo Belda, Malgorzata Broncel, Gwendolin Fuchs, David Anaguano, Stephanie D. Nofal, Moritz Treeck

## Abstract

Fever is a hallmark of malaria. Several studies have linked febrile temperatures to reduced parasite viability, but also to increased cytoadhesion, a key driver of pathology. However, different mechanisms have been proposed to cause changes in cytoadhesion and parasite sensitivity to heat. Here, we demonstrate that exposure of *Plasmodium falciparum*-infected red blood cells (iRBCs) to physiologically relevant febrile heat stress (39 °C), derived from patient data, enhances cytoadhesion through increased trafficking of the major virulence factor PfEMP1 to the iRBC surface. This phenomenon is not limited to PfEMP1 and common laboratory strains, as it extends to the surface nutrient channel PSAC in four clinical isolates of diverse geographic origin. The increased surface protein display occurs without changes in overall protein expression or parasite developmental progression. Using phosphoproteomics and proximity labelling, we find that elevated temperature also increases trafficking and phosphorylation of exported proteins into the RBC. Enhanced export is likely reliant on the presence of a transmembrane domain as shown by NanoLuc reporter assays. Collectively, our results indicate that febrile temperatures commonly experienced during infection can accelerate protein export, likely at the parasitophorous vacuole. This enhanced export following heat stress is relevant because increased cytoadhesion could influence disease severity through earlier iRBC sequestration and elevated bound parasite mass.

## Introduction

Malaria remains one of the most devastating infectious diseases and is caused by unicellular eukaryotic parasites of the *Plasmodium* genus. Of the six species that infect humans, *Plasmodium falciparum* is responsible for the vast majority of malaria-related deaths. An estimated 249 million malaria cases and 608,000 deaths occur annually ^1^. The pathology of malaria arises during the asexual replication phase of *P. falciparum* within red blood cells (RBCs). This cycle involves RBC invasion, intracellular growth, and the production of daughter merozoites, which are released upon host cell rupture, leading to exponential parasite growth.

Fever is a hallmark of malaria and a defining feature of the host immune response to *P. falciparum* infection. Malaria-associated fever is thought to arise from pathogen-associated and damage-associated molecular patterns (PAMPs and DAMPs) released during RBC rupture as parasites egress ^2–4^. Febrile heat stress is frequently encountered by *P. falciparum*, and the parasite has likely evolved mechanisms to adapt to and prosper under these conditions. As asexual parasites progress through the 48-hour life cycle, they develop through three key stages: ring stage (0–24 hours post-invasion (hpi)), trophozoite stage (24-40 hpi), and schizont stage (40-48 hpi).

Previous studies found that the first half of the asexual life cycle (0-24 hpi) is more resistant to heat stress than the second half (24-48 hpi) ^5^. Research into the impact of elevated temperatures on *P. falciparum* growth and gene regulation has provided key insights into the parasite’s response to heat stress. Random mutagenesis screening of 992 *P. falciparum* genes at 41 °C identified more than 200 mutants with significantly altered growth responses ^6^. Transcriptomic analyses of wild-type *P. falciparum* following heat stress (40 °C and 41 °C) revealed significant changes in gene expression. Oakley et al. ^7^ found that these changes mirror apoptotic cell responses, while Tinto-Font et al. ^8^ identified a transcription factor (PfAP2-HS) required for parasite growth at both normal (37 °C) and elevated (40 °C) temperatures.

Cytoadhesion of *P. falciparum*-infected red blood cells to the vascular endothelium is a key virulence trait. It is mediated by the exported and surface-displayed Erythrocyte Membrane Protein 1 (PfEMP1), which prevents splenic clearance. The specific PfEMP1 variant expressed determines the site of sequestration. For example, expression of the VAR2CSA variant enables infected red blood cells to bind chondroitin sulfate A (CSA) in the placenta ^9^. PfEMP1-mediated sequestration in host organs is unique to *P. falciparum* and is a major driver of disease severity, contributing to serious complications such as cerebral and placental malaria ^10^. Fever-like temperatures have been reported to increase PfEMP1 on the surface of RBCs, although this has been contested as described in more detail below^11^.

Approximately 10% of *P. falciparum* encoded proteins are predicted to be exported beyond the parasite plasma membrane, with many thought to pass through the parasitophorous vacuole lumen (PVL) and parasitophorous vacuole membrane (PVM) into the RBC cytosol, although the exact routes for transmembrane proteins are not fully understood ^12–14^. These exported proteins remodel the host cell, mediating processes such as nutrient acquisition and immune evasion. Exported proteins contain one of two trafficking elements: PEXEL-containing proteins, which undergo N-terminal cleavage before export, and PEXEL-negative exported proteins (PNEPs) which are not cleaved but contain transmembrane domains required for export ^13,15,16^.

PfEMP1 is trafficked to the red blood cell surface via parasite-derived membranous organelles known as Maurer’s clefts ^17–20^. This process relies on several exported proteins, many unique to *P. falciparum*, including FIKK4.1, an exported kinase that localises to the red blood cell periphery ^21^.Deletion of FIKK4.1 leads to a 55% reduction in PfEMP1-mediated cytoadhesion ^21^. Even though several proteins have been identified that are important for PfEMP1 trafficking, the precise mechanisms that underlie PfEMP1 surface translocation remain poorly understood ^19,22^.

Several studies have examined the effects of febrile heat stress on iRBC cytoadhesion, showing contradictory results. Udomsangpetch *et al.* ^11^ reported that heat stress increased iRBC cytoadhesion to endothelium expressing the PfEMP1 cognate receptor, correlating with higher PfEMP1 levels on the iRBC surface. In contrast, Oakley *et al.* ^7^ found no change in the proportion of iRBCs displaying PfEMP1 on the surface following heat stress. Furthermore, trophozoite stage iRBCs become more rigid under heat stress, a property predicted to aid parasite sequestration in organs ^23^. Zhang et al. ^24^ reported an alternative mechanism whereby cytoadhesion increased due to elevated phosphatidylserine (PS) and not PfEMP1 on the iRBC surface following heat stress. One protein that has been linked to both heat stress and PfEMP1 surface levels is the *P. falciparum* exported protein HSP70x. Charnaud et al. ^25^, but not Cobb et al. ^26^, found HSP70x to be important for PfEMP1 trafficking, although both studies concluded that HSP70x is dispensable for intraerythrocytic parasite growth at 37 °C. Charnaud *et al.* ^25^, further reported no significant growth differences in HSP70x knockout parasites under heat stress or nutrient-limiting conditions. In contrast, Day et al. ^27^ observed reduced parasitaemia in HSP70x knockdown parasites in the cycle following heat stress, compared to wild-type strains. In summary, whether febrile temperatures modulate cytoadhesion and by what mechanism remains unclear. Given that increased cytoadherent iRBC biomass in febrile patients could contribute to disease severity, further studies are important.

The divergent outcomes of the above-mentioned studies could be due to differences in the heat-stress conditions used, including those that affect parasite viability. Furthermore, *in vitro* culture of *P. falciparum* can cause gene loss of exported proteins, and transfections can impose a genetic bottleneck that may enrich such mutants ^28^. It is therefore not uncommon to observe defects in cytoadhesion after generating transgenic parasites. Consequently, loss of cytoadhesion without restoring the phenotype by gene complementation or using conditional approaches should be interpreted with caution.

Here, we assessed how febrile heat stress influences exported protein trafficking and cytoadhesion in *P. falciparum.* We first defined three key criteria: 1) a physiologically relevant heat stress commonly occurring in malaria patients, 2) its impact on parasite growth across asexual stages, and 3) the normal timing of PfEMP1 trafficking to the iRBC surface. Using this framework, we tested whether febrile heat stress modulates iRBC cytoadhesion and its underlying mechanism. We show that heat stress increases cytoadhesion of iRBCs, coinciding with a greater proportion of infected cells displaying PfEMP1 on their surface. Comparative phosphoproteomic analysis and proximity labelling of the Maurer’s clefts protein environment suggest that elevated temperatures enhance protein trafficking into the red blood cell cytosol. NanoLuciferase (NanoLuc) fusion proteins and compartment-specific isolation confirmed a greater abundance of PfEMP1 in the RBC cytosol following heat stress. Constitutively expressed reporter proteins suggest that transmembrane domains are key contributors to this increased trafficking, independent of overall protein expression levels. This increased surface trafficking is not limited to PfEMP1. It also extends to the Plasmodial Surface Anion Channel (PSAC), a parasite-derived nutrient channel, across four *P. falciparum* clinical isolates from diverse geographic origins.

## Results

### A physiologically relevant and non-destructive febrile heat stress increases iRBC cytoadhesion and surface PfEMP1 display

While a variety of temperatures mimicking febrile heat stress have been used in previous studies, these have generally not been assessed in relation to parasite viability, or common febrile temperatures in patients. Therefore, we sought to initially establish conditions that reflect the typical febrile response in malaria patients, focusing on a time window when PfEMP1 is exported and parasite viability remains unaffected.

To approximate the most common febrile temperature in individuals infected with *P. falciparum*, we analysed patient temperature data from two publicly available supplementary datasets ^29,30^. The average tympanic febrile temperature (defined as greater than 38 °C) in *P. falciparum* malaria cases was 38.9 °C and 39.1 °C, respectively (Figures 1A-B). Temperatures ≥ 41 °C were rarely recorded in either dataset. Analysis of microarray data from Mok *et al.* ^29^ showed that higher patient temperatures were associated with a younger average age of circulating parasites, possibly reflecting increased cytoadherence and sequestration of more mature stages during fever (Figure 1C).

**Figure 1.**
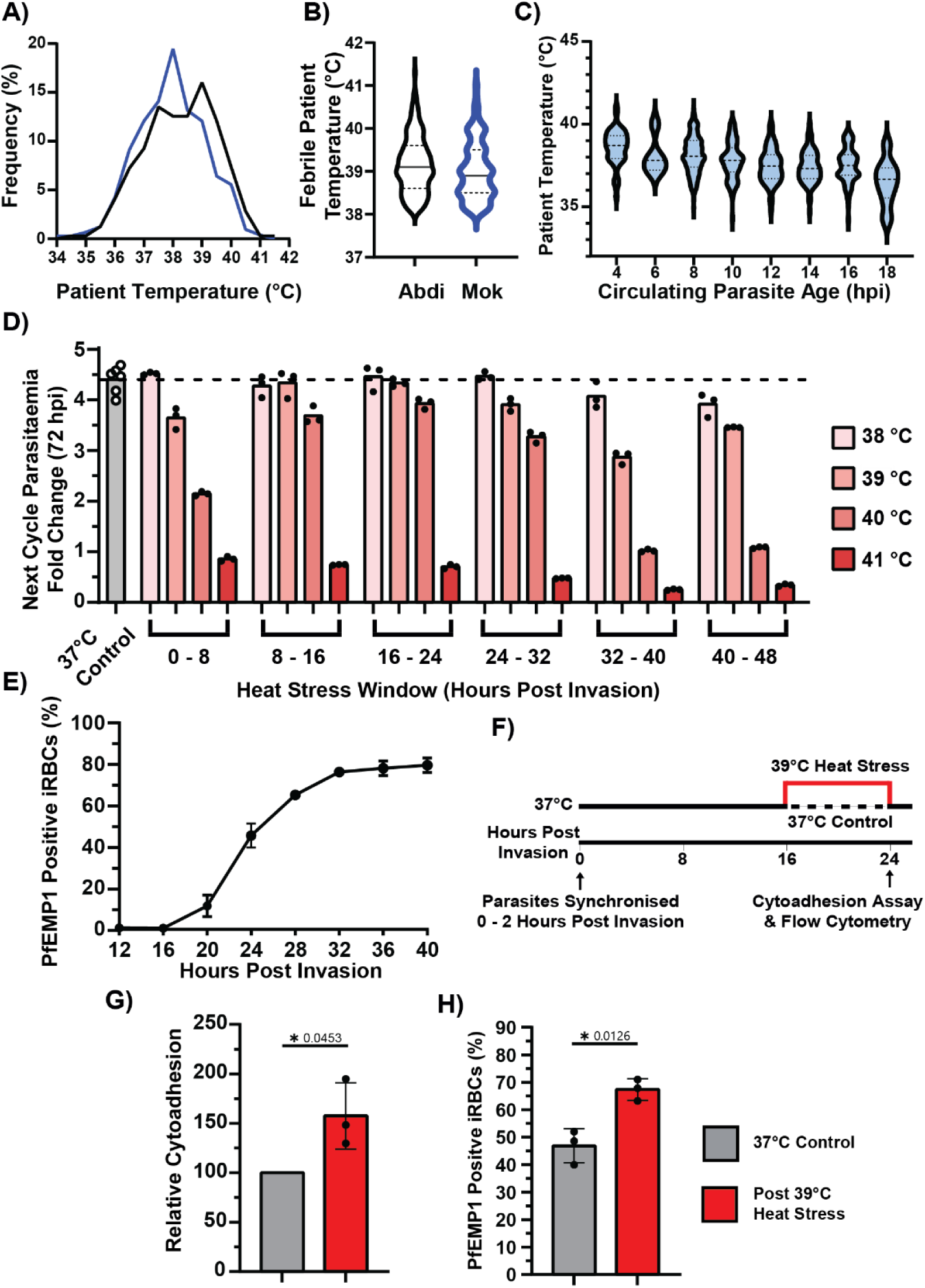
A physiologically relevant, non-destructive febrile heat stress increases the percentage of *P. falciparum* infected red blood cells expressing PfEMP1 on their surface and enhances cytoadhesion to the cognate receptor. (A) Relative distribution of patient body temperature from two studies (Mok *et al.* ^29^ and Abdi *et al.* ^30^) that included accessible temperature data from *P. falciparum* infected patients (n = 1043 (blue) and n = 827 (black), respectively). (B) Average febrile temperature, defined as >38 °C, was 38.9 °C and 39.1 °C in Mok *et al.*, and Abdi *et al.*, respectively. (C) Estimated average age of circulating parasites from Mok *et al.* microarray data plotted against recorded patient temperature. (D) Synchronised infected red blood cells (iRBCs) were exposed to heat stress for different 8-hour windows during the first 48 hours post invasion (hpi). When not heat-stressed, parasites were cultured under normal conditions at 37°C. Parasitaemia was measured by flow cytometry in the following cycle at 72 hpi. N=3 (technical triplicates) for heat-stressed cultures; n=6 (biological replicates) for 37°C control cultures. Dashed lines represent the average next cycle parasitaemia fold change of control cultures. (E) Percentage of iRBC positive for PfEMP1 (VAR2CSA) surface expression from 12 to 40 hpi when cultured normally at 37 °C. Flow cytometry gating strategy is shown in Figure 1 – Supplement 2. (F) Schematic representation of the heat stress application to synchronous NF54 DiCre parasites. (G) Following heat stress, more iRBCs bound to CSA under flow (1 dyne/cm^2^). (H) Significantly more iRBCs were positive for PfEMP1 (VAR2CSA) after heat stress. For G and H n=3 biological replicates. Error bars displayed are ± 1 SD. Statistical significance was determined using unpaired t-tests of log-transformed data with Welch’s correction. Flow cytometry and cytoadhesion assays were performed at room temperature.

To assess the impact of heat stress on *P. falciparum* (NF54 DiCre strain ^31^) growth and replication, parasites were exposed to elevated temperatures (38-41 °C) for eight hours across six different developmental time windows. Parasitaemia was measured in the subsequent cycle (Figure 1D). Similar to the observations by Kwiatkowski ^5^, we found that parasites in the first half of the cycle (0-24 hpi) were more resistant to heat stress than those in the second half (24-48 hpi). However, our data further reveal that recently invaded parasites (0-8 hpi) are less resistant to elevated temperatures compared to later ring-stage parasites (8-16 hpi and 16-24 hpi). We also observe that exposure to 41 °C for eight hours is highly destructive to the parasite, irrespective of the developmental stage at which it is applied.

To test whether PfEMP1 surface translocation is affected at the mean febrile temperature observed in patients, we first determined the percentage of iRBCs maintained at 37 °C that display PfEMP1 (VAR2CSA) on the surface from 12-40 hpi. These experiments revealed a dynamic window of surface expression between 16-32 hpi, consistent with previous reports ^32,33^ (Figure 1E). These foundational data informed the identification of a heat stress window (39 °C) that mimics fever-range temperatures, overlaps with VAR2CSA trafficking to the iRBC surface (16-24 hpi), and was not expected to impair parasite growth.

We next subjected tightly synchronised *P. falciparum* cultures to elevated temperatures (39 °C) or control conditions (37 °C) (Figure 1F) to assess the impact of heat on cytoadhesion and PfEMP1 surface translocation. Following heat stress, significantly more iRBCs (57.6% ±19.4% more than non-heat stressed controls) cytoadhered under physiological flow conditions (1 dyne/cm²) compared to those maintained at 37 °C, which was set as the baseline and normalised to 100% (Figure 1G). This increase in cytoadhesion correlated with a significantly higher percentage of iRBCs displaying PfEMP1 (VAR2CSA) on their surface (67.4% ± 3.93% versus 46.9% ± 6.18%) compared to the control (Figure 1H). Importantly, heat stress between 16-24 hours post-invasion did not accelerate parasite development through the asexual life cycle (Figure 1 – Supplement 1).

Collectively, these data, in line with Udomsangpetch *et al.* ^11^ show that more iRBCs cytoadhere following febrile heat stress as the parasites develop from ring stages to trophozoites. This coincides with a significantly greater number of iRBCs displaying PfEMP1 on the surface. The mechanism underlying the increased translocation of PfEMP1 to the surface remains unknown.

### Red blood cells infected with four *P. falciparum* strains from different geographic origins exhibit increased sensitivity to sorbitol following heat stress

To investigate whether proteins other than PfEMP1 are trafficked differently during heat stress, we sought to identify another cell surface protein that is both detectable and essential for *in vitro* growth, ensuring its expression across all *P. falciparum* strains. One such protein complex is the Plasmodial Surface Anion Channel (PSAC), which facilitates nutrient uptake and is exported to the surface of iRBCs. PSAC comprises three proteins: RhopH2, RhopH3, and one of several cytoadherence-linked asexual gene (CLAG) variants ^34,35^, which contain a putative transmembrane domain (TMD) ^36–38^. All three components are expressed during the late stages of the parasite life cycle. These pre-formed proteins are stored in the parasitophorous vacuole membrane (PVM) following invasion and are trafficked into the host cell via the *Plasmodium* translocon of exported proteins (PTEX) complex at a time similar to PfEMP1 ^14,39,40^.

Unlike PfEMP1, PSAC-mediated nutrient uptake is essential for parasite survival under *in vitro* conditions ^42–44^. Functional PSAC on the iRBC surface transports the small molecule sorbitol into the red blood cell cytosol, disrupting the osmotic balance of the cell and rupturing the iRBC (Figures 2A-B). Consequently, the percentage of ruptured iRBCs following sorbitol treatment serves as a surrogate marker for functional PSAC present at the iRBC surface ^45^.

**Figure 2.**
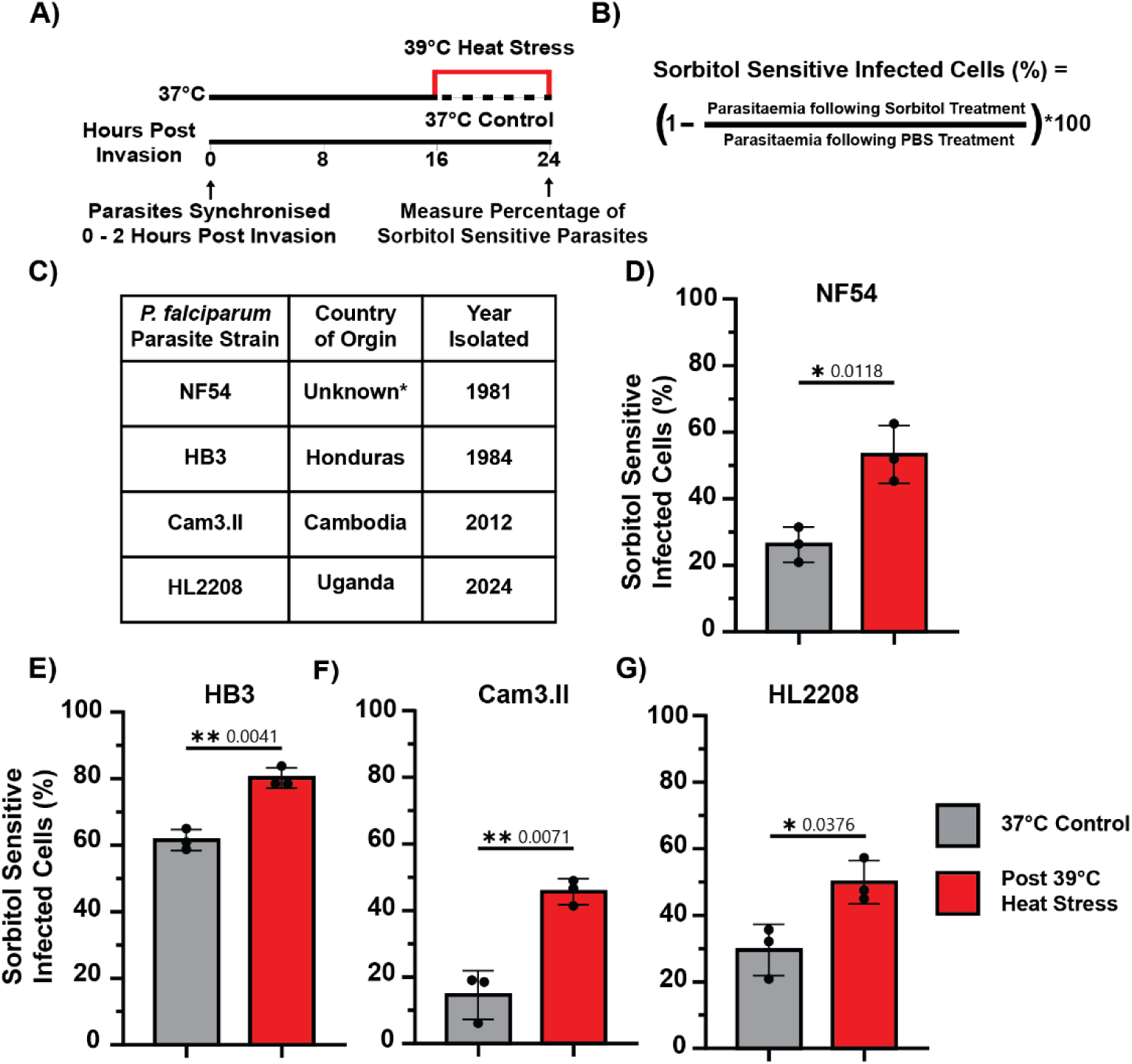
Heat stress increases sorbitol sensitivity in red blood cells infected with four *P. falciparum* strains from diverse geographic origins. (A) At 24 hpi following an eight-hour heat stress (39 °C), parasite cultures were treated with PBS and sorbitol, and the parasitaemia was determined and compared to that of a 37 °C control. Sorbitol sensitivity is conferred by a functional nutrient channel (PSAC) trafficked by the parasite onto the surface of iRBCs. (B) As sorbitol sensitive parasites rupture during treatment, the percentage of sorbitol sensitive parasites is calculated by comparing parasitaemia following sorbitol treatment to parasitaemia following PBS treatment. (C) Geographic origins of *P. falciparum* isolates (*NF54 is predicted to have an African origin but was isolated in the Netherlands ^41^). (D–G) After heat stress, NF54 (D), HB3 (E), Cam3.II (F), and HL2208 (G) strains exhibited significantly higher sensitivity to sorbitol treatment. Data represents three biological replicates (N=3). Statistical significance was assessed using unpaired t-tests on log-transformed data with Welch’s correction. All assays were performed at room temperature.

We found that significantly more NF54 iRBCs were sensitive to sorbitol treatment following heat stress (53.3% ±5.2%) compared to normal conditions (26.2% ±8.6%) (Figures 2C-D). Sorbitol lysis affected only iRBCs and was blocked by the PSAC inhibitor furosemide, showing that heat stress does not impact uninfected RBC integrity or induce PSAC independent sorbitol uptake (Figure 2 – Supplement 1). To assess whether heat-induced changes in protein surface translocation are conserved across *P. falciparum* strains, we extended our analysis to three geographically distinct isolates: HB3 (Honduras, 1984), Cam3.II (Cambodia, 2012), and HL2208 (Uganda, 2024) ^46–49^. In all three strains, heat stress (39 °C) resulted in a significant increase in iRBC sensitivity to sorbitol lysis (Figures 2E-G), consistent with findings in the NF54 strain. Unless PSAC components are modified (for example through post-translational modifications) to increase PSAC activity following heat stress, these results indicate that earlier trafficking of PSAC to the iRBC surface under febrile heat stress is a conserved phenomenon across diverse *P. falciparum* isolates. Culturing parasites in sub-lethal furosemide concentrations or in reduced nutrient media lead to reduced parasitaemia (Figure 2 – Supplement 2). However, the parasitaemia is not further reduced following heat stress. This shows that increased PSAC levels/activity do not enhance parasite survival under conditions of limited nutrient availability either from furosemide-induced nutrient deprivation or a reduced nutrient media composition. Collectively, these results indicate that the increased membrane trafficking described above is not restricted to PfEMP1.

### A *P. falciparum* heat stress phosphoproteome reveals substantial enrichment in phosphorylation of exported proteins by exported FIKK kinases

We next sought to identify factors that influence protein trafficking to the iRBC surface. Our previous work identified the exported protein kinase FIKK4.1 as important for efficient surface translocation of PfEMP1 ^21^. This kinase is also required for phosphorylation of RhopH3 (PF3D7_0905400), a component of the PSAC complex ^21,50^ and may therefore play a role in regulating the trafficking of both proteins. FIKK4.1 is part of the FIKK kinase family, which includes 19 kinases in *P. falciparum*, 18 of which are likely exported into the RBC. Several other exported kinases, including FIKK10.2, a kinase localised to the Maurer’s clefts, phosphorylate proteins involved in PfEMP1 export but do not affect PfEMP1 trafficking at 37 °C ^50,51^. To broadly investigate whether FIKK kinases respond to elevated temperatures and potentially contribute to increased surface translocation of PfEMP1 and PSAC, we measured the phosphoproteome of iRBCs following heat stress and control conditions.

Following the same heat stress window previously shown to increase cytoadhesion (16-24 hpi, 39 °C), parasites were returned to 37 °C until 30 hpi. At this point, iRBCs were enriched from uninfected cells by density separation and lysed in 8 M urea. Proteins were digested and labelled with tandem mass tags (TMT), and enriched phosphopeptides were identified by LC-MS/MS (Figure 3A, Supplementary Tables 1-2). A total of 196 phosphopeptides (1.2% of those detected) were differentially phosphorylated following heat stress, with 143 showing increased phosphorylation (>1 Log2FC) and 53 showing decreased phosphorylation (<-1 Log2FC). Notably, 71% of the heat-enriched phosphopeptides mapped to exported proteins, which far exceeds the approximately 10% of the total proteome predicted to be exported (Figures 3B-C) ^12,13^. Analysis of non-phosphorylated peptide levels showed no major changes in either the human or parasite proteomes following heat stress (Figure 3 – Supplement 1). This indicates that neither parasite development nor total protein expression account for the observed increase in surface PfEMP1.

**Figure 3.**
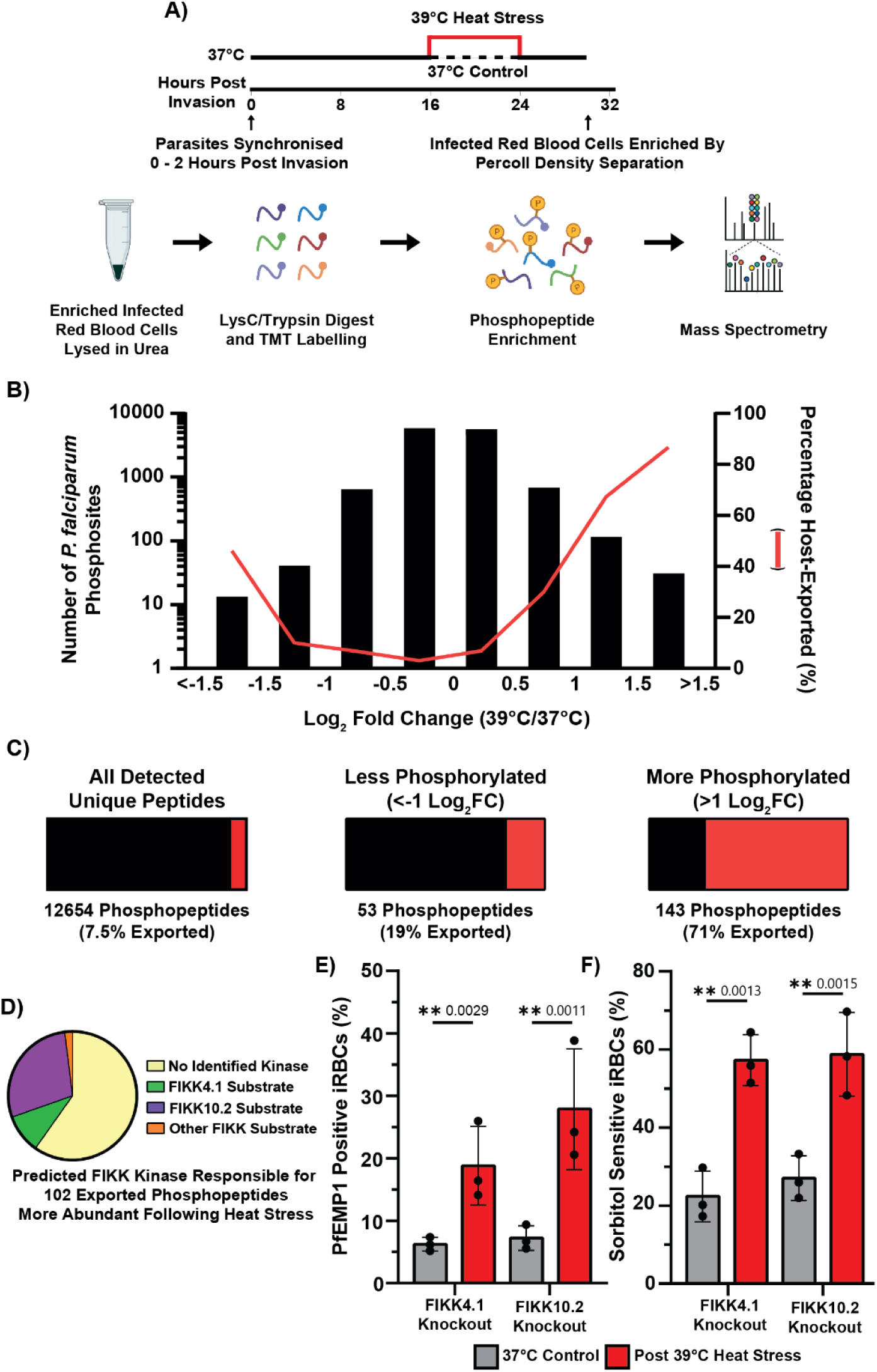
Febrile heat stress results in increased phosphorylation of host cell exported proteins. (A) Overview of infected red blood cell lysate preparation for phosphoproteomics. (B) Histogram of 12654 unique *P. falciparum* phosphopeptides, showing enrichment of phosphopeptides from exported parasite proteins following heat stress. (C) Percentage of unique phosphopeptides that are from parasite exported proteins (red). Following heat stress the significantly more abundant phosphopeptides (>1 Log2FC) are enriched for exported proteins (71%) compared to the 12564 phosphopeptides not changing (7.5%). N=2 Biological Replicates. (D) Comparing the 102 exported phosphopeptides more abundant following heat stress against confident FIKK substrates identified FIKK10.2 and FIKK4.1 as likely major contributors to heat stress phosphorylation. The remaining phosphopeptides had not previously been linked to a specific kinase (No Identified Kinase) by Davies et al. ^52^. Neither FIKK4.1 nor FIKK10.2 are required for the heat stress-induced increase in PfEMP1 surface expression (E) or for the increased sorbitol sensitivity of iRBCs following heat stress (F), as shown using synchronised kinase conditional knockout (RAP-treated) parasite lines. N=3 biological replicates. Error bars displayed are ± 1 SD. Statistical significance was determined using the one-way ANOVA test with the Benjamini, Krieger and Yekutieli FDR correction. Antibody staining, flow cytometry and sorbitol treatment was performed at room temperature. For (E) and (F) parasites were heat stressed between 16-24 hpi at 39 °C, sorbitol treatment and antibody staining were performed at 24 hpi.

We have previously mapped the phosphorylation events that depend on each FIKK kinase under normal temperature (37 °C) conditions ^21^. To determine whether the heat-induced phosphosites were novel or dependent on known FIKKs, we compared heat-induced phosphosites with our prior datasets ^21^. This analysis suggested that most heat-dependent phosphorylation could be largely attributed to FIKK10.2 and, to a lesser extent, FIKK4.1 (Figure 3D). To validate these results, we performed a second phosphoproteomic analysis following heat stress conditions using several previously generated exported FIKK knockout strains (FIKK1, FIKK4.1, FIKK4.2, all FIKK9s, FIKK10.1, and FIKK10.2). This confirmed that FIKK10.2 is required for the majority of phosphorylation increases observed in exported proteins during heat stress (Figure 3 – Supplement 2, Supplementary Table 3).

Interestingly, while FIKK4.1 was previously shown to be necessary for efficient PfEMP1 surface translocation at 37 °C ^21^, we still observed a significant increase in PfEMP1 surface levels in the absence of FIKK4.1 following heat stress. Despite being critical for most heat-stress phosphorylation events, FIKK10.2 was also not required for the heat-induced increase in PfEMP1 surface presentation or sorbitol sensitivity (Figures 3E-F).

To test whether disruption of proteins targeted by FIKK kinases during heat stress might help identify required mediators of early trafficking, two additional DiCre-mediated conditional knockout strains of HA-tagged lines were generated. HSP70x (PF3D7_0831700), previously implicated to have a role in PfEMP1 trafficking and heat stress survival, showed increased phosphorylation under heat stress conditions ^25,27^. PF3D7_0702500, a Maurer’s cleft-residing protein ^53,54^ exhibited the highest number of phosphosites with increased phosphorylation in response to heat stress. Both tagged proteins localised as previously reported and were efficiently excised upon RAP treatment (Figure 3 – Supplements 3-6) ^25,26,53^. However, neither HSP70x nor PF3D7_0702500 disruption was found to have an effect on heat stress survival, PfEMP1 trafficking, PSAC surface expression at 37 °C, or were required for accelerated heat stress trafficking (Figure 3 – Supplements 7-12).

Collectively, these data show that elevated temperature leads to increased phosphorylation of exported proteins, with the majority of this modification mediated by the Maurer’s cleft-localising kinase FIKK10.2. However, neither FIKK4.1, FIKK10.2 nor the putative substrates HSP70x and PF3D7_0702500 are required for the enhanced trafficking of PfEMP1 or the increased sorbitol sensitivity, which is likely driven by elevated PSAC trafficking. Based on these findings, we reasoned that the observed increase in phosphorylation may result from greater protein flux into or through subcellular compartments where FIKK4.1 and FIKK10.2 localise.

### The FIKK10.2-TurboID Maurer’s cleft proxiome during normal and febrile temperatures

FIKK10.2 is localised to the Maurer’s clefts, which are key trafficking hubs established by the parasite in the RBC. To measure trafficking of proteins through the Maurer’s Clefts, we generated a C-terminal fusion of endogenous FIKK10.2 and TurboID (FIKK10.2-TurboID) (Figure 4 – Supplement 1). In the presence of biotin and ATP, TurboID generates and releases reactive biotin molecules that bind to lysine residues in proximal (up to ∼10 nm) proteins ^55^. Biotinylated proteins are then digested, and the resulting biotinylated peptides are enriched by immunoprecipitation, quantified by LC-MS/MS and mapped to the corresponding proteins ^50^.

To assess how the FIKK10.2-TurboID proxiome changes during heat stress, we measured three conditions: (A) continuous biotin labelling without heat stress to identify proteins that are either persistently or transiently proximal to FIKK10.2 throughout the asexual cycle, (B) biotin labelling from 16-24 hpi without heat stress to capture interactions during PfEMP1 trafficking, and (C) biotin labelling with heat stress from 16-24 hpi to determine heat-induced changes in FIKK10.2’s local protein environment (Figures 4A-B).

**Figure 4.**
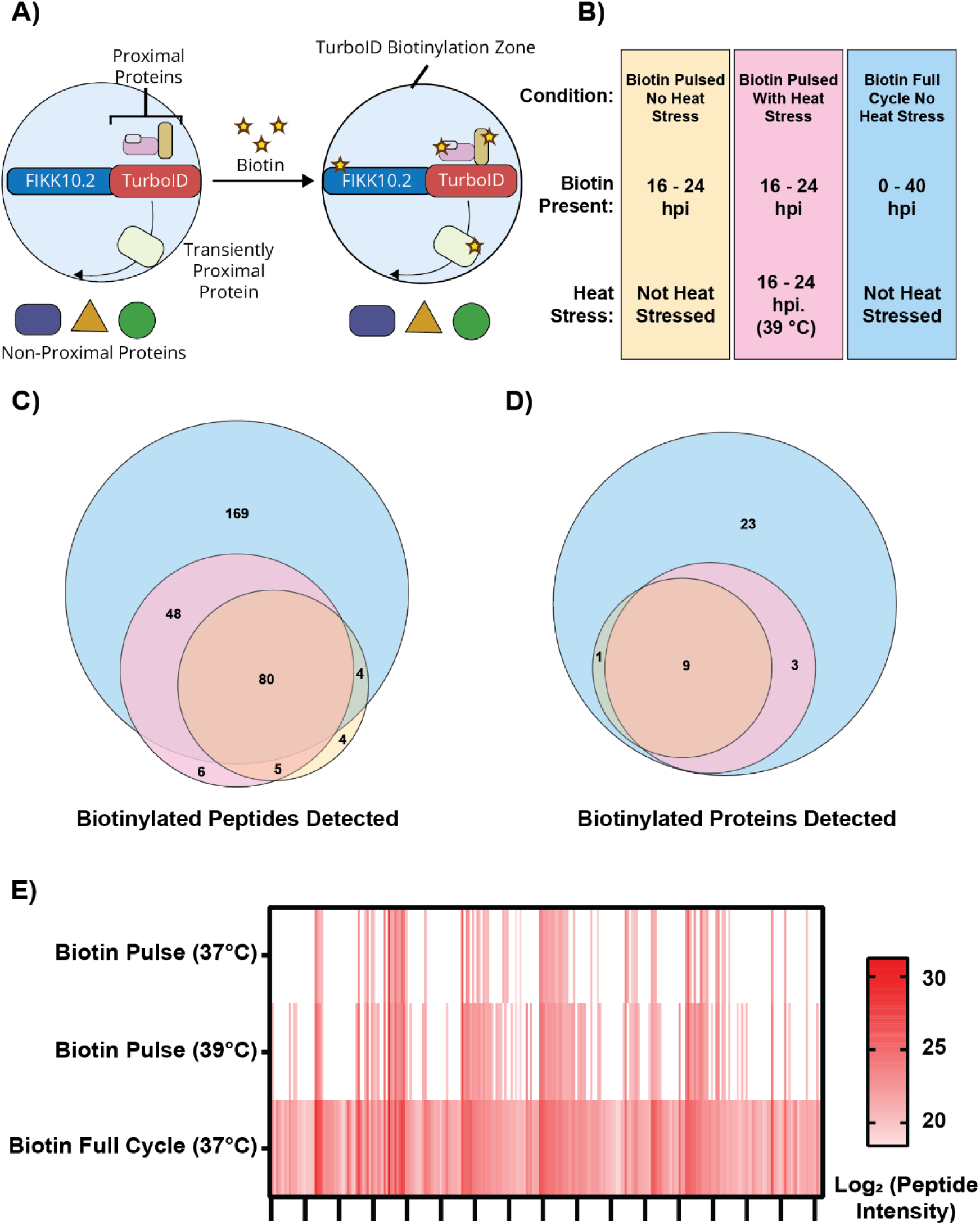
FIKK10.2-TurboID reveals changes to the Maurer’s cleft protein environment during heat stress. (A) In the presence of biotin, transgenic parasites expressing FIKK10.2, a Maurer’s cleft residing protein, fused to TurboID were used to biotinylate proximal proteins. (B) FIKK10.2-TurboID parasites were cultured under three different conditions: biotin was either present throughout the life cycle, or added as an eight-hour pulse (16–24 hpi) in the presence or absence of heat stress at 39 °C. At 40 hpi, late-stage iRBCs were enriched by Percoll density separation and lysed. Proteins were subsequently digested, and biotinylated peptides were enriched and identified. (C) Proportional Venn diagram of detected peptides across the three conditions. (D) Proportional Venn diagram of detected proteins in each condition. (E) Heat map of biotinylated peptide intensity across the three tested conditions. N = 3 biological replicates. For (C) and (D), valid peptides were defined as those detected in at least two replicates, and valid proteins were defined as those with at least two different valid peptides detected.

Across all conditions, a total of 316 unique biotinylated peptides were detected, comprising 278 from *P. falciparum* and 38 from human proteins. In condition A, where biotin was present throughout the parasite life cycle, 256 *P. falciparum* peptides were consistently detected in every replicate. Of these, 231 peptides (90%) mapped to 39 exported proteins, underscoring the specificity and robustness of this peptide-level enrichment approach ^50^. These proteins are very likely to be either trafficked through or localised to the Maurer’s clefts (Supplementary Tables 4-5). Many of these proteins have predicted transmembrane domains and by mapping the FIKK10.2-TurboID enriched peptides on their projected topology, the red blood cell cytosol facing domain can be predicted (Figure 4 – Supplement 2).

Although six peptides were detected only in the 39 °C heat-stressed biotin-pulsed condition (Figure 4C), they originate from proteins also present in the other conditions (Figures 4D-E). Thus, these TurboID data indicate that FIKK10.2 does not encounter unique proteins during heat stress that are absent under normal conditions. Based on these data, we hypothesised that a greater number of exported proteins are in proximity to FIKK10.2 during febrile heat stress, suggesting increased trafficking of proteins to or through the Maurer’s clefts. These proteins are phosphorylated by FIKK10.2 during their transit. One possible origin of increased trafficking of exported proteins could be increased translocation of proteins through the PVM into the RBC cytosol.

### Febrile heat stress increases PfEMP1 (VAR2CSA) trafficking into the RBC cytosol and enhances its surface presentation on a greater proportion of iRBCs

To directly assess whether PfEMP1 is trafficked earlier to the red blood cell surface during heat stress, we endogenously tagged the VAR2CSA variant with NanoLuc (Figure 5A, Figure 5 – Supplement Figure 1) ^56,57^. Following heat stress, a significantly greater proportion of iRBCs displayed surface-localised PfEMP1 tagged with NanoLuc, as detected by α-VAR2CSA staining and flow cytometry, indicating that the fusion protein retains its capacity to traffic into the host cell and reach the iRBC surface (Figure 5B).

**Figure 5.**
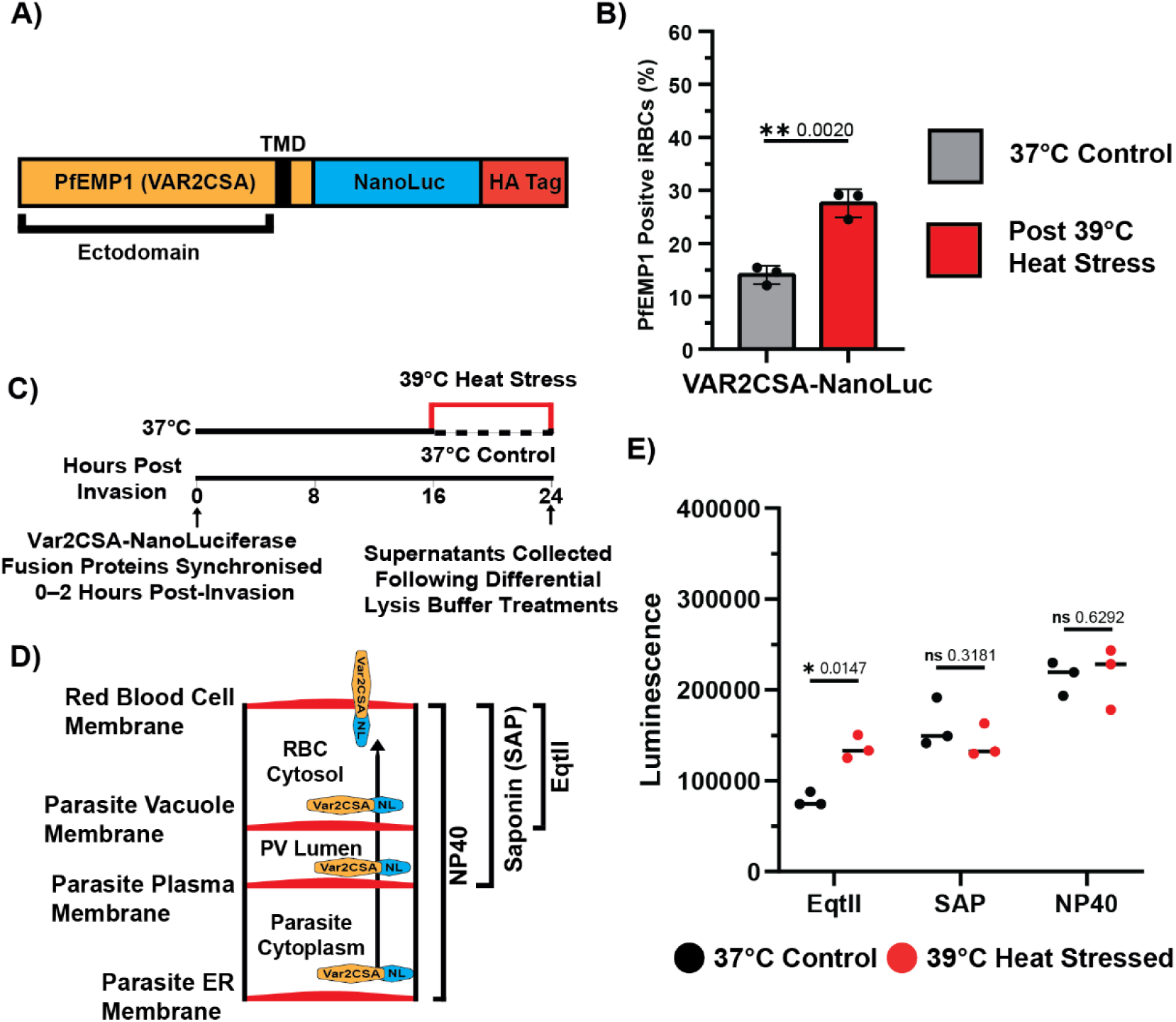
Febrile heat stress enhances trafficking of endogenously tagged PfEMP1 (VAR2CSA) to the RBC cytosol and surface. (A) A transgenic *P. falciparum* strain expressing NanoLuc fused to the C-terminus of VAR2CSA was generated. (B) Following heat stress, a greater proportion of VAR2CSA-NanoLuc expressing parasites displayed PfEMP1 on the iRBC surface compared to parasites maintained at 37 °C. N = 3 biological replicates. Error bars represent ±1 SD. Statistical significance was determined using an unpaired t-test on log-transformed data with Welch’s correction. (C) At 24 hpi, following heat stress or continuous culture at 37 °C, cells were differentially permeabilised using three treatments. NanoLuc activity was measured in the supernatants. (D) Overview of the compartments released by each permeabilisation treatment: EqtII permeabilises the RBC membrane while preserving the PVM, SAP releases proteins from the RBC cytosol and PV lumen, and NP40 solubilises all non-nuclear proteins from the parasite, PV, and RBC cytosol. (E) Significantly higher NanoLuc activity was detected in the supernatant of heat-stressed parasites only following EqtII permeabilisation. Assays were performed at room temperature. Data represent the mean of three biological replicates, each averaged from three technical replicates. Statistical significance was determined using the one-way ANOVA test with the Benjamini, Krieger and Yekutieli FDR correction.

To quantify export efficiency under normal (37 °C) and heat-stressed (39 °C) conditions, we measured NanoLuc activity across three distinct compartments: (A) the RBC cytosol, (B) the parasitophorous vacuole (PV), and (C) the total parasite/RBC lysate. These compartments were selectively isolated in the supernatant following different permeabilisation protocols. Equinatoxin II (EqtII) permeabilises the RBC membrane while leaving the PVM intact, saponin (SAP) releases proteins from both the RBC cytosol and PV lumen ^58^, and NP40 detergent solubilises all non-nuclear proteins within the parasite, PV, and RBC cytosol (Figures 5C-D).

Importantly, NanoLuc activity in the NP40 supernatant did not differ significantly between control and heat-stressed conditions, suggesting that total VAR2CSA expression remains unchanged (Figure 5E). This supports our hypothesis that *P. falciparum* does not markedly accelerate its developmental program or significantly alter protein expression in response to heat stress, and that increased cytoadhesion under febrile conditions is more likely due to altered protein trafficking.

Notably, following heat stress, NanoLuc activity significantly increased (58% more) in the EqtII permeabilised supernatant but not in the SAP fraction (Figure 5E). This suggests that heat stress promotes trafficking of VAR2CSA from the PV into downstream compartments such as the Maurer’s clefts and ultimately to the RBC surface.

### Transmembrane domain containing exported reporter proteins are trafficked more efficiently into the red blood cell cytosol during febrile heat stress

To evaluate whether the enhanced trafficking of exported proteins under heat stress (39 °C) is a general phenotype or restricted to transmembrane domain-containing proteins such as PfEMP1 and the PSAC component CLAG, we generated four NanoLuc reporter constructs with variable N-terminal protein fusions (Figure 6 – Supplement 1). All four reporters were expressed under the constitutive HSP90 (PF3D7_0708400) promoter and integrated into the EBA165 pseudogene locus (PF3D7_0424300).

To first test whether other TMD-containing proteins are differentially trafficked into the RBC during febrile heat stress, the export of PF3D7_0702500-NanoLuc was measured. PF3D7_0702500 is a Maurer’s cleft-resident protein ^53,54^, which we found to be significantly more phosphorylated and more biotinylated by FIKK10.2-TurboID following heat stress (Supplementary Tables 1 and 4). The trafficking of PF3D7_0702500-NanoLuc into the iRBC cytosol mirrored that observed for VAR2CSA-NanoLuc, with PF3D7_0702500-NanoLuc being moderately more abundant (30%) in the EqtII fraction following heat stress (39 °C) compared to normal conditions (37 °C) (Figure 6A-C).

**Figure 6.**
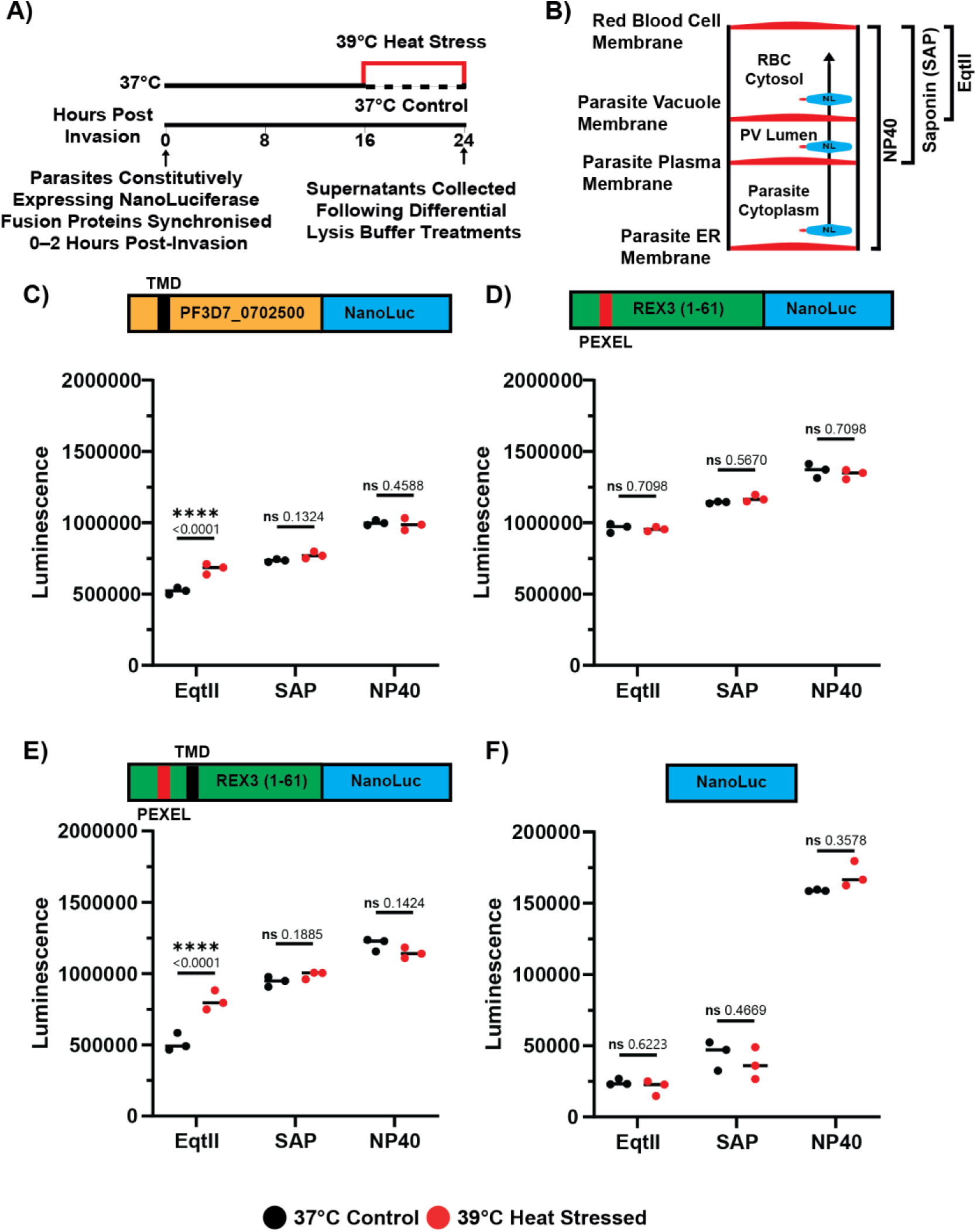
Constitutively expressed exported NanoLuc strains identify transmembrane domains as key determinants for increased export into the host cell during febrile temperatures. (A) Tightly synchronised *P. falciparum* parasites constitutively expressing one of four NanoLuc reporter constructs were exposed to heat stress (39 °C) or maintained at 37 °C between 16–24 hpi. At 24 hpi, cells were subjected to three different differential permeabilisation treatments. NanoLuc activity in the resulting supernatants was measured. (B) Schematic representation of the cellular compartments released by each permeabilisation method. All assays were performed at room temperature. (C) The constitutively expressed PF3D7_0702500-NanoLuc reporter, but not D) the REX3-NanoLuc reporter, showed significantly increased NanoLuc activity in the EqtII-permeabilised supernatant following heat stress. REX3-NanoLuc contains the first 61 amino acids of REX3, which are sufficient to mediate export ^59,60^. (E) Fusion of the PF3D7_0702500 TMD to the REX3-NanoLuc reporter reduced luminescence in the EqtII-permeabilised supernatant under non-stress conditions. However, increased luminescence was observed following heat stress. (F) Reporters lacking an N-terminal fusion to an exported *P. falciparum* protein showed negligible luminescence in the supernatant following SAP or EqtII treatment. Across all four constructs, no significant differences in luminescence were observed between SAP and NP40-permeabilised samples when comparing conditions with and without heat stress. Data represent the mean of three biological replicates, each averaged from three technical replicates. Statistical significance was determined using the one-way ANOVA test with the Benjamini, Krieger and Yekutieli FDR correction.

As a non-TMD-containing protein reporter, we generated REX3-NanoLuc, which contains the first 61 amino acids of REX3 that are sufficient for export ^59,60^. This reporter did not show a significant increase in abundance in any of the three permeabilisation fractions following heat stress (Figure 6D).

These data suggest that TMD containing proteins are specifically affected in trafficking under heat conditions. To directly test this, the TMD from PF3D7_0702500 was added to REX3-NanoLuc to generate REX3-TMD-NanoLuc. Trafficking of REX3-TMD-NanoLuc into the RBC cytosol was less efficient than REX3-NanoLuc at 37 °C. However, following heat stress, 57.6% more REX3-TMD-NanoLuc was detected in the RBC cytosolic fraction compared to normal temperature conditions (Figure 6E). A non-exported NanoLuc strain was also generated as a negative control and showed negligible signal in the exported compartments, with no significant change under heat stress (Figure 6F).

Collectively, these data suggest that elevated febrile temperatures alleviate a trafficking bottleneck between the PV and the RBC cytosol, at least for the TMD-containing proteins examined in this study. Notably, addition of the PF3D7_0702500’s TMD to a soluble reporter was sufficient to enhance export under heat stress, indicating that this may represent a broader mechanism by which TMDs facilitate temperature-responsive trafficking into the RBC cytosol.

## Discussion

The mechanism underlying increased iRBC cytoadhesion following febrile heat stress is not well understood, with both PfEMP1-dependent and independent pathways proposed. Variability in experimental heat stress conditions, including differences in temperature that affect parasite viability, as well as differences in duration and timing within the asexual cycle, likely contributes to the divergent findings reported across studies. We show that elevated temperatures at multiple stages of the parasite life cycle negatively affect parasite viability. These growth-impairing conditions are likely to trigger significant stress or apoptosis-like responses. To minimise the potential for such stress responses to confound cytoadhesion outcomes, we selected a non-destructive heat stress condition (39 °C, 16-24 hpi) for our experiments. More extreme protocols may not accurately reflect the febrile temperatures typically experienced by malaria patients. In addition to the PfEMP1-dependent mechanism examined here, increased phosphatidylserine (PS) on the surface of iRBCs following 1 hour at 40 °C has been observed and coincides with increased cytoadhesion to CHO cells expressing endothelial receptors ^24^. Although a role for PfEMP1 in this increased cytoadhesion was not identified previously ^24^, our findings indicate that PfEMP1 surface expression contributes possibly alongside PS to the heightened cytoadhesion during fever. However, genetic deletion of proteins required for PS exposure and PfEMP1 surface translocation may be necessary to unequivocally establish the individual contribution of each to overall cytoadhesion.

Our data confirm a previous study reporting increased PfEMP1 on the red blood cell surface following febrile heat stress ^11^. We propose that this increase in surface trafficking results from a reduction in a trafficking bottleneck during elevated temperature. Elevated temperature does not only affect PfEMP1 surface translocation but also enhances trafficking of other TMD-containing proteins into the RBC. This likely results in increased PSAC levels at the iRBC surface, although we cannot fully exclude the possibility that the observed increase in sorbitol sensitivity is due to enhanced PSAC activity following heat stress, for example through post-translational modifications. However, we consider this explanation less likely, as deletion of the kinases known to phosphorylate PSAC components had no measurable effect on sorbitol sensitivity. Using several complementary approaches, including NanoLuc reporter assays, proteomics, DNA content measurements, egress timing, and assessments of subsequent replication, we conclude that under our heat stress conditions (39 °C, 16 to 24 hpi), parasite development does not accelerate. These findings indicate that the observed phenotypes are due to differences in protein trafficking.

While heat stress induces changes in the Maurer’s cleft proxiome and phosphoproteome, we did not identify any proteins that were altered exclusively under heat stress compared to 37 °C. Later in the asexual cycle, heat stressed iRBCs reach levels of PfEMP1 surface expression comparable to those maintained at 37 °C (Figure 3 – Supplements 8 and 11). This suggests that proteins involved in accelerated trafficking during heat stress are also present and likely functional at normal temperatures, rather than being uniquely expressed or specifically localised in response to elevated temperature. However, it is worthwhile to note that some phosphorylation sites increase upon heat stress that do not depend on FIKK10.2 nor FIKK4.1. Several of these sites could also not be attributed to previously identified FIKK substrates, indicating that some may be mediated by FIKKs other than FIKK4.1 and FIKK10.2, or by host cell kinases.

The mechanism by which proteins are differentially exported at elevated temperatures remains unknown, although differential trafficking requirements for TMD and non-TMD-containing proteins have been proposed at both the PPM and PVM ^61,62^. Our reporter assay measures export into the RBC compartment but cannot definitively distinguish whether the trafficking differences occur at the level of the PVM or later, for example between the Maurer’s clefts and the red blood cell plasma membrane. Nevertheless, the PTEX complex, which mediates protein translocation from the PV through the PVM into the RBC cytosol ^14^, is a likely candidate to facilitate increased export of TMD-containing proteins under heat stress. We excluded HSP70x to be involved in protein trafficking as we show that conditional deletion of HSP70x does not affect PfEMP1 surface translocation.

What may be the consequences of accelerated PfEMP1 trafficking, or protein surface export during fever in a human infection? Our data show that a greater proportion of iRBCs are sensitive to sorbitol following heat stress, suggesting that more cells may present functional PSAC at the surface. Increased function of PSAC could potentially enhance nutrient uptake under these strained conditions. Elevated PfEMP1 levels at the iRBC surface during heat stress may help counteract the negative effects of increased cellular rigidity that occur at higher temperatures ^24^. As cytoadhesion is thought to prevent filtration of the rigidified iRBC in the spleen ^63^, concurrent increased cytoadhesion may need to occur with increased stiffness. How altered protein trafficking during heat stress affects the interplay between iRBC stiffness and their potential to sequester and obstruct small vessels remains to be addressed.

If increased body temperature leads to an increased cytoadhering parasite load, malaria symptoms may be exacerbated. The observed reduction in the age of circulating parasites at higher temperatures suggests that cytoadhesion may occur earlier under febrile conditions. An alternative explanation is that the younger parasite age reflects a recent, synchronised egress event, which has been proposed to trigger fever ^3^. However, tightly synchronised parasite replication is likely rare in fever-inducing natural infections.

Quantifying the synchronicity of natural *P. falciparum* infections is inherently difficult, primarily due to the sequestration of mature parasite stages via cytoadherence and the ethical concerns of continuous monitoring without treatment. Controlled human malaria infection (CHMI) studies, such as Kapulu et al. ^64^, sampled circulating parasites twice daily using quantitative PCR. Infections were allowed to progress for up to 21 days or until fever symptoms develop. Even in these tightly controlled conditions, circulating parasites are often detectable at every 12-hour interval, suggesting that they are not tightly synchronized. As parasite burden increases, complete absence of circulating parasites becomes increasingly rare, further indicating asynchronous development. Polyclonal infections, resulting from repeated mosquito bites and the introduction of genetically distinct parasite strains, also appear to drive asynchronicity ^65^. This is particularly relevant in high-transmission settings where individuals may receive, on average, more than 20 Anopheles mosquito bites per day ^66^. Importantly, *P. falciparum* fever should not be conflated with synchronicity. As noted by Kitchen in Malariology (1949) ^67^, the febrile patterns of *P. falciparum* are so diverse that the term ’‘atypical’’ may not meaningfully be applied. While fever may influence parasite dynamics, it is unlikely that only specific stages experience it; all stages are plausibly exposed to host fever responses during an infection.

Ultimately, it is unclear if febrile conditions promote survival of the parasite or the host, it is likely that a sustained fever (>39 °C, >8 hours) restricts parasite proliferation. However, the frequency of sustained high malarial fevers is not well documented. Moreover, data investigating parasite resilience to febrile heat stress are based mainly on lab-adapted parasites, which are not under selection pressure to survive elevated temperatures. Antipyretic drugs reduce fever, but whether they should be used during viral and bacterial infections is uncertain ^68^. This ambiguity also extends to malaria where a meta-analysis of several clinical studies investigating malaria patient outcome following antipyretic treatment found that there was insufficient data to confirm or refute an impact of antipyretic measures on parasitaemia or malarial illness ^69^. However, a recent clinical trial showed a positive correlation of antipyretic treatment with host survival in Malaria infection indicating further studies in this area are warranted ^70^. Furthermore, a recent pre-print demonstrates increased binding of iRBCs to blood vessels using 3D models as a result of fever conditions ^71^. Together, increased PfEMP1 on the surface earlier with increased host-cell binding would exacerbate the negative effects on fever on host-survival if the overall impact of fever on parasite survival is not taken into account. To directly test the impact of fever on parasite survival versus cytoadhesion, well controlled studies in human populations will be required in the future.

## Acknowledgments

We thank members of the Treeck lab for critical discussions and additionally Dr. Christian Gnann, Dr. Franziska Hildebrandt and Miss Ana Matias for their feedback on this manuscript. We would like to thank members of the Sateriale, Blackman and Bernabeu lab groups for critical discussions throughout the project. We also thank the Crick Science Technology Platforms (Proteomics, Flow Cytometry, Media Preparation and Light Microscopy) for their outstanding technical support. Special thanks to PlasmoDB for providing a critical resource. We thank Colin Sutherland for providing the HL2208 *P. falciparum strain*, Lars Hviid for the α-VAR2CSA (PAM7.5) antibody, Tobias Spielmann for the α-SBP1 antibody, and the European Malaria Reagent Repository for the α-KAHRP antibody.

M.T. is supported by funding from the ERC (Grant Number 10144428), which also provided funding to H.B., G.F. and D.A., The Francis Crick Institute (Grant Numbers CC2132 & CR2023/030/2132), which also provided funding to H.B. The Francis Crick Institute Science Technology Platforms (Grant Number CC0199) and the FCT (Fundación para a Ciencia e Tecnologia (Grant Number 2023.06167.CEECIND)). S.D.N. is funded by an Early-Career Award Wellcome Trust grant (225686/Z/22/Z). G.F. is funded by the FCT through a PhD fellowship (Grant Number 2024.02695.BD). The Francis Crick Institute and its Science Technology platforms receive core funding from Cancer Research UK, the UK Medical Research Council and the Wellcome Trust.

## Author Contributions

D.J. designed and performed phenotypic analysis. D.J. and M.T. conceived the study. D.J. and M.B. processed the proteomic samples. D.J.,H.B., and M.B. analysed the proteomic data. D.J., H.B., G.F., S.D.N., and D.A. performed parasite genetic manipulations. G.F. performed the immunofluorescent microscopy and D.A. carried out sorbitol lysis validation experiments. D.J. and M.T. wrote the original manuscript. All authors were involved in critically reviewing and editing the manuscript.

## Materials and Methods

### *In vitro* maintenance and synchronisation of parasites

Human erythrocytes infected with asexual stages of *P. falciparum* were cultured at 37°C in complete medium. Complete medium was prepared using 1 L RPMI-1640, supplemented with 5 g AlbuMAX II (ThermoFisher Scientific), 0.292 g L-glutamine, 0.05 g hypoxanthine, 2.3 g sodium bicarbonate, 0.025 g gentamicin and 5.957 g HEPES. The washed and purified red blood cells were provided by the UK Blood and Transfusion Service. Parasites were cultured following standard procedures in a gas atmosphere of 90% N2, 5% CO2, and 5% O2 ^72^. Parasite cultures were synchronised using Percoll (GE Healthcare) to isolate mature schizont-stage parasites. The purified schizonts were incubated at 37°C in complete medium with fresh RBCs for 2-4 hours in a shaking incubator. A second Percoll purification was performed to remove any remaining schizonts, and the remaining cells were resuspended in 5% sorbitol for 10 minutes at 37°C before being washed into complete medium. These two steps resulted in highly synchronous parasites with minimal debris in the solution.

### Generation of transgenic *P. falciparum* strains

*P. falciparum* transfections were performed as described before ^73,74^. Transgenic Plasmodium falciparum NF54::DiCre ^31^ parasite lines were generated using CRISPR-Cas9. Unless otherwise specified, the NF54::DiCre parental strain was used as a wild type-like reference throughout this manuscript. Suitable guide RNAs (gRNAs) were identified using the Eukaryotic Pathogen CRISPR guide RNA/DNA Design Tool ^75^. Complementary oligonucleotides corresponding to the 19 nucleotides nearest the selected PAM sequence were synthesised (Integrated DNA Technologies, IDT), phosphorylated using T4 polynucleotide kinase, annealed and ligated into the pDC_Cas9_hDHFRyFCU ^76^ vector digested with BbsI. To produce compatible sticky ends between the annealed oligonucleotides and the BbsI-digested vector, the forward oligonucleotide included a 5′-ATTG overhang, and the reverse oligonucleotide included a 5′-AAAC overhang. gRNAs targeting the gene of interest were assembled using appropriate oligonucleotide pairs. Repair templates were designed in three different configurations. For conditional knockouts, templates included LoxP-containing introns flanking a recodonised, HA-tagged gene followed by a T2A skip peptide and a drug selection cassette^77^. For TurboID fusions, templates encoded a recodonised gene C-terminally fused to TurboID with a V5 tag, followed by a T2A skip peptide and drug selection marker. For NanoLuc reporters, the construct comprised the HSP90 promoter driving expression of a variable N-terminal domain fused to NanoLuciferase, followed by a T2A skip peptide and a selection cassette ^74^. Gene blocks encoding recodonised genes, TurboID, and NanoLuciferase were synthesised (IDT). All repair templates were flanked by gene-specific homology arms and included restriction sites to enable excision of the linear repair templates. Prior to transfection 20 μg of a Cas9-expressing guide plasmid and 20 μg of a linearized repair template plasmid, were ethanol precipitated. Ethanol precipitation was performed by adding 0.1 volume of 3 M Sodium Acetate (pH 5.2) and 2 volumes of 100% ethanol, followed by at least 2 hours incubation at -20 °C. DNA was subsequently centrifuged at 4 °C for 30 minutes and washed twice with 70% ethanol. DNA pellets were air-dried and resuspended in 10 μl Tris-EDTA (TE) buffer. Synchronised egressing schizonts (NF54 DiCre) were enriched by 60% Percoll before washing in complete medium. The resuspended DNA was mixed with 90 μl P3 Primary cell solution (Lonza) and was used to resuspend egressing schizonts before transfer to transfection cuvette.

Transfection was performed by electroporation using the FP158 programme of an Amaxa 4D Electroporator machine (Lonza). Following transfection, the parasites were transferred to pre-warmed flasks containing 2 ml complete medium and 300 μl fresh uRBCs. After 40 minutes of gentle shaking incubator at 37 °C, 8 ml complete medium were added to each flask. Transfected parasites were cultured without selection for 24 hours, then selection was first performed with 2.5 nM WR99210 (Jacobus Pharmaceuticals) for 48 hours. After WR99210 resistant parasites emerged, they were further selected with 225 μg/ml G418 (Gibco). Correct integration of the repair template into the parasite’s DNA was confirmed by overlapping PCR of the 5’ and 3’ regions (Supplementary Table 6).

### Conditional knockout of DiCre expressing parasites

To induce DiCre-mediated *LoxP* site recombination, synchronised ring-stage parasites were treated with either 100 nM RAP (Sigma) or an equivalent volume of DMSO to act as a control. After four hours at 37 °C, iRBCs were washed twice with RPMI and returned to culture in complete medium. Samples for DNA and protein extraction were taken at least one replication cycle (48 hours) after RAP treatment.

### PfEMP1 (VAR2CSA) cell-surface detection

50 μL of iRBC culture was centrifuged (3000 g, 1 minute) and pelleted cells were blocked in 200 μL 3% BSA in PBS for 15 minutes at room temperature. Cultures were then resuspended in 50 μL of 1% BSA in PBS with 1:200 α-VAR2CSA ^78^ (PAM7.5) human monoclonal antibody (a kind gift from Lars Hviid) and incubated for 30 minutes at 37 °C. Cells were then pelleted and washed in 1 mL of PBS three times to remove unbound primary antibody. Cultures were then resuspended in 50 μL of 1% BSA in PBS with 1:200 α-Human PE-conjugated secondary antibody (12-4998-82, ThermoFisher), 1:2000 Hoechst (ThermoFisher) and incubated for 30 minutes at 37 °C. Cells were then pelleted and washed in 1 mL of PBS three times to remove unbound secondary antibody. Labelled cells were then measured by flow cytometry (FACSVerse, BD Biosciences) without fixation.

### Application of heat stress *to P. falciparum* cultures

To heat stress parasites, separate cell culture incubators were set to different temperatures. Correct temperature was determined independently of the incubator screen display by multiple probes. When required, flasks were moved from one incubator to another to heat stress or to return to 37 °C after heat stress. An effort was made to not keep flasks out of the incubator for long and flask transfer took less than 30 seconds. When heat stress was being applied the opening of the incubators was kept to a minimum.

To measure the temperature dynamics of medium within heat stressed flasks, a temperature logging probe (Fisherbrand™ 90009-36) was inserted into the media, and the temperature was recorded every 30 seconds (Accuracy: ±0.5 °C, Resolution: 0.1 °C).

### Infected red blood cell cytoadhesion assay

Cytoadhesion assays were performed as described previously ^50^. 100 μl of CSA (Sigma-Aldrich) (1 mg/ml in PBS) was added to each channel of an untreated Ibidi μ-slide VI_0.4_ and left to incubate overnight at 4 °C. The channels were blocked with 3% BSA in PBS for one hour at room temperature before three washes in PBS and one final wash in RPMI without Albumax. Parasites were synchronised to a two-hour invasion window (10% parasitaemia, 1% haematocrit). At 24 and 40 hpi, 500 μL of iRBC culture was resuspended in 4 mL of RPMI without Albumax and drawn through an Ibidi μ-slide VI_0.4_ previously coated with CSA and blocked in BSA. Using a Harvard syringe pump (Sigma) 1 dyne/cm^2^ (0.1 Pa) shear stress was applied across each channel (0.55 mL/minute). At the same pressure and flow rate the channels were then washed with 4 mL of RPMI without Albumax containing Hoechst DNA stain (1:2000, ThermoFisher). Ten frames (40x) across the centre of the microchannel were taken with Nikon Eclipse Ti2 Microscope. Cytoadherent cells were quantified via ImageJ imaging software as Hoechst positive cells. Multiple conditions were measured in parallel, and binding was normalised to the 37 °C control.

### P. falciparum growth assays

To assess parasite growth, cultures were synchronised to a two-hour invasion window and parasitaemia was adjusted to 0.5-1%. Parasitaemia was typically measured in the initial cycle (cycle 1) and the subsequent two cycles using flow cytometry. At 24 (cycle 1), 72 (cycle 2), and 120 (cycle 3) hours post-invasion (hpi), a 50 µL aliquot of iRBC culture (1-4% haematocrit) was mixed thoroughly with 50 µL of 2× glutaraldehyde (GA) (0.4%) in a 96-well plate. Plates were sealed and stored at 4 °C. Once all timepoints were collected, 50 µL was removed from each fixed well and combined with 50 µL of 2× SYBR Green DNA stain (1:5000, ThermoFisher) in PBS, then incubated at 37 °C for 30 minutes. Following incubation, 100 µL of PBS was added to each well, and the plate was analysed using a high-throughput plate attachment on the FACSVerse flow cytometer (BD Biosciences). Infected cells were gated based on SYBR Green fluorescence intensity, and parasitaemia was calculated for each timepoint.

### Infected red blood cell sorbitol sensitivity assay

Parasites were synchronised to a two-hour invasion window (1–10% parasitaemia and 1–5% haematocrit). To assess sorbitol sensitivity of iRBCs, two 200 µL aliquots of iRBC culture were taken from each sample and centrifuged at 3000 × *g* for 2 minutes. The cell pellets were washed with 1 mL of PBS and centrifuged again. Each aliquot was then resuspended in either 5% sorbitol or PBS, with both solutions pre-warmed to 37 °C. Following a 10-minute incubation at 37 °C, cells were washed four additional times with 1 mL of PBS. Pelleted cells were either fixed and parasitaemia determined by SYBR Green staining (1:10,000, ThermoFisher),or analysed directly without fixation. Flow cytometry was performed as described previously using a FACSVerse flow cytometer (BD Biosciences). The percentage of sorbitol-sensitive parasites was calculated using the following formula: 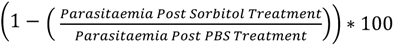. To assess whether sorbitol mediated lysis was restricted to infected red blood cells and inhibited by the PSAC inhibitor furosemide following heat stress both non heat stressed and heat stressed parasite cultures were treated with 200 µM furosemide for 10 min at 37 °C. Cultures were maintained at 2% haematocrit with parasitaemia greater than 10%. Following pre-treatment with furosemide cultures were resuspended in 5% sorbitol in PBS with 200 µM furosemide maintained. After a 10 min incubation samples were centrifuged and haemoglobin release was quantified by measuring the absorbance of the supernatant at 415 nm using a plate reader. As additional controls, cultures were incubated in PBS alone without sorbitol or treated with saponin to achieve complete cell lysis. Supernatants from saponin treated samples were diluted five-fold to ensure measurements fell within the linear range of detection.

### *P. falciparum* growth following heat stress when cultured in the presence of furosemide or with reduced nutrient medium

NF54 DiCre parasites were synchronised to a two-hour invasion window and 1% parasitaemia before replacing the medium with either complete medium (as described above), complete medium with 150 µM furosemide or reduced nutrient medium. The composition of the reduced nutrient media was from Molina et al. ^79^ and is detailed in Supplementary Table 7. A DMSO vehicle control was added to the complete medium and reduced nutrient medium. Between 16 and 24 hpi, infected red blood cells were either heat stressed at 40 °C or maintained at 37 °C and parasitaemia was determined in the subsequent cycle at 72 hpi by flow cytometry. Following heat stress, cultures were maintained at 37 °C.

### NanoLuciferase *P. falciparum* export assay

NanoLuciferase (NanoLuc) expressing transgenic parasites were synchronised to a two-hour invasion window (1-3% parasitaemia, 1% haematocrit). To gather samples to measure activity, a 600 μL aliquot of parasite culture was pelleted (3000 × g, 1 minute) and resuspended in 1 mL of PBS. This solution was then further divided into three 325 μL aliquots. These three aliquots were then pelleted and the supernatant removed. The pellets were either resuspended in 200 μL of 0.1% NP40 in PBS, 0.03% Saponin (Sigma) in PBS or three haematolytic units (HUs) of Equinatoxin II (a kind gift from Mike Blackman) in PBS. These solutions were then incubated at 37 °C for 10 minutes. Following incubation, the samples were centrifuged (10000 × g for 5 minutes) and 100 μL of the supernatant was collected without disturbing the pellet. The buffer composition of each of the fractions was normalised by the addition of 100 μL of the other two buffers. Therefore, each condition had a final volume of 300 μL and contained 1 HU of Equinatoxin II, 0.01% Saponin and 0.033% NP40 in PBS. These fractions were then kept at -20 °C until required.

To measure NanoLuc activity, fractions were first thawed on ice. 100 μL was transferred into a white opaque 96 well plate (Corning). Technical triplicates were typically measured in three different wells. NanoGlo substrate provided in the Nano-Glo® Luciferase Assay System (Promega) was diluted 1:500 in PBS. 100 μL of the substrate dilution was then added to each well for a final substrate concentration of 1:1000. Luminescence was then measured with 225 gain in a Cytation 96 well plate reader (Agilent, BioTek).

### Immunofluorescence assay

Thin blood smears of iRBC cultures on microscope slides were submerged in ice-cold methanol at −20 °C to fix the cells. Slides were then wrapped in lint-free tissue and stored with silica beads at -80 °C until required. For labelling, slides were thawed at 37 °C in the presence of silica beads for 20 minutes. Once dry, hydrophobic rings were drawn on the slides, one for each labelling condition. Slides were blocked with 3% BSA in PBS for 1 hour at room temperature. Cells were labelled with primary antibodies diluted in PBS with 1% BSA for at least 1 hour at room temperature, applying 25 μL of antibody solution to each hydrophobic ring. After three washes with PBS, secondary antibodies containing BSA and DAPI were added and incubated under the same conditions for 1 hour. Finally, slides were washed, mounted with ProLong Gold antifade reagent (ThermoFisher, P36930) and covered with a coverslip. Primary antibodies included rat α-HA (3F10, Merck (11867423001), 1:250), Rabbit α-SBP1 ^80^ (a kind gift from Tobias Spielmann, 1:5000) and Mouse α-KAHRP (The European Malaria Reagent Repository,mAb 18.2, 1:000). After three final washes with PBS, coverslips were mounted over the stained cells with Prolong Gold antifade reagent (Invitrogen) containing DAPI and sealed with nail polish. Images were taken using a Ti-E Nikon microscope using a x100 TIRF objective at room temperature equipped with a LED-illumination. Alternatively, Higher magnification images were acquired on a VisiTech instant SIM (VTChapteriSIM) microscope using a 150x oil-immersion objective and 1.5 μm z-axis steps. Image processing was performed in FIJI.

### Immunoblotting

Schizonts were enriched by Percoll density separation and lysed in 1% SDS RIPA buffer supplemented with 1 mM DTT. Soluble cell extracts were first separated by SDS-PAGE before transferring onto a nitrocellulose membrane (Biorad) by Transblot Turbo. As a loading control, VersaBlot™ CF680nm (Biotium) was added to protein samples before SDS PAGE to enable visualisation of total protein content on the subsequent blot. Membranes were blocked overnight at 4 °C in 2% BSA, 0.2% Tween-20 in PBS. After blocking membranes were probed with primary antibody for 1 hour at room temperature, before three washes in 0.2% Tween-20 in PBS (PBS-T). Primary antibodies included rat α-HA (3F10, Merck (11867423001), 1:3000) and Streptavidin-HRP (Pierce (21130), 1:5000). Washed membranes were then probed with the appropriate secondary antibody HRP conjugate (1:10000, Abcam) for an hour at room temperature. Membranes were finally washed with PBS-T three more times before adding HRP substrate and buffer (Immobilon ECL Ultra, Sigma) was added and the blot imaged by ChemiDoc (BioRad). The total protein stain was visualised at 680 nm.

### Sample preparation for phosphoproteomic profiling of *P. falciparum*

The protocol for generating and processing *P. falciparum* phosphoproteome samples was previously described in detail by Davies *et al.* ^21^. Synchronised NF54 *P. falciparum* parasites (0–2 hpi) were cultured at 37 °C and subjected to either heat stress (39 °C from 16–24 hpi) or maintained at 37 °C. While not heat stressed, parasites were kept at 37 °C. Infected red blood cells (iRBCs) were enriched at 30 hpi using a 60% Percoll density gradient centrifugation and washed three times with PBS. Parasite pellets were lysed in 8 M urea in 50 mM HEPES (pH 8.5), supplemented with protease inhibitors (cOmplete™ Mini, EDTA-free, Roche) and phosphatase inhibitors (PhosSTOP™, Roche). Lysates were sonicated (3 × 30 seconds, 30% duty cycle, on ice), clarified by centrifugation (13,000 × g, 30 min, 4 °C), and the supernatants were snap-frozen in liquid nitrogen and stored at −80 °C.

For phosphoproteomic analysis, a TMT 10-plex (Thermo Fisher Scientific) was used for wild-type parasites ± heat stress (two biological replicates per condition, using red blood cells from independent donors). Two additional channels were used for uninfected red blood cells ± heat stress (N = 1 each), and the remaining four channels were used for unrelated experiments. A second experiment used TMTpro 16-plex (Thermo Fisher Scientific) labelling across seven knockout lines (FIKK1, FIKK4.1, FIKK4.2, FIKK9s, FIKK10.1, FIKK10.2) ± heat stress, alongside wild-type and uninfected RBC controls (N=1 per condition). These knockout strains were previously generated and validated by Davies *et al.* ^21^. For the conditional knockout strains, gene disruption was induced by DiCre-mediated excision as described above, in the preceding intraerythrocytic cycle to ensure full loss of gene function. These strains, alongside the NF54 wild-type, were then resynchronised at the start of the cycle (0-2 hpi) prior to heat stress and sample collection.

### Reduction, alkylation, and protein digest of phosphoproteome samples

Once thawed, protein concentration was determined by a BCA protein assay (Pierce). 1 mg of each lysate was subsequently reduced with 10 mM DTT for 60 minutes at room temperature and alkylated in the dark with 20 mM iodoacetamide for 30 minutes at room temperature. Excess iodoacetamide was then quenched with 10 mM DTT. Lysates were diluted with 50 mM HEPES pH 8.5 to < 2 M urea and each digested with 5 µg of LysC (WAKO) for 2-3 hours at 37 °C and then overnight with 30 µg of trypsin (Thermo Scientific) at 37 °C.

### Sep-Pak desalting

The samples were acidified with trifluoroacetic acid (TFA) (Thermo Fisher Scientific) to a final concentration of 0.4% (v/v) and left on ice for 10 minutes. All insoluble material was removed by centrifugation (1,600 × g, 10 minutes, 4 °C), and the supernatants were desalted using Sep-Pak Lite C18 cartridges (Waters™) in conjunction with a vacuum manifold. Columns were initially washed with 3 mL of acetonitrile, conditioned with 1 mL of 50% acetonitrile and 0.5% acetic acid in H₂O, and then equilibrated with 3 mL of 0.1% TFA in H₂O. The acidified samples were loaded, desalted with 3 mL of 0.1% TFA in H₂O, washed with 1 mL of 0.5% acetic acid in H₂O, and finally eluted with 1.2 mL of 50% acetonitrile and 0.5% acetic acid in H₂O. Each sample was subsequently dried by vacuum centrifugation.

### TMT labelling

Samples were dissolved in 1 mL of 50mM Na-HEPES pH 8.5 and 30% anhydrous acetonitrile (v/v) and labelled with the respective TMT reagents (Thermo Fischer Scientific). Both TMT-10plex (2.4mg reagent/1mg sample) and TMT-16plex (3mg reagent/1mg sample) were used in two separate experiments. Labelling reactions were carried out for one hour at room temperature followed by quenching with 0.3 % hydroxylamine for 15 minutes at room temperature and sample acidification (pH ∼2) with formic acid. After verification of labelling efficiency via mass spectrometry, the lysates were mixed in a 1:1 ratio, vacuum dried and desalted on Sep-Pak C18 cartridges.

### Phosphopeptide enrichment and high pH sample fractionation

Phosphopeptides were sequentially enriched using the High Select SMOAC protocol. In the first step a High Select TiO2 phosphopeptide enrichment kit (Thermo Fisher Scientific) was used according to the manufacturer instructions.

The supernatant from the initial TiO₂ enrichment was vacuum dried and a second enrichment was performed using the High Select Fe-NTA phosphopeptide enrichment kit (Thermo Fisher Scientific), following the manufacturer’s instructions. The flow-through, containing non-phosphorylated peptides (representing the total proteome), was retained and stored at −80 °C. Phosphopeptide eluates from TiO₂ and Fe-NTA enrichment were pooled and fractionated using the Pierce High pH Reversed-Phase Peptide Fractionation Kit (Thermo Fisher Scientific), following the manufacturer’s instructions for unlabelled native peptides. Resulting 8 fractions were subsequently dried under vacuum to prepare for LC-MS/MS.

### LC-MS/MS (phosphoproteomics) and data analysis

TMT-10plex: samples were resuspended in 0.1% TFA and loaded onto a 50 cm EasySpray PepMap column (75 μm inner diameter, 2 μm particle size; Thermo Fisher Scientific) fitted with an integrated electrospray emitter. Reverse-phase chromatography was performed using an RSLC nano U3000 system (Thermo Fisher Scientific) with a binary buffer system: solvent A consisted of 0.1% formic acid and 5% dimethyl sulfoxide (DMSO) in water, while solvent B contained 80% acetonitrile, 0.1% formic acid, and 5% DMSO. Chromatographic separation was carried out at a flow rate of 275 nL/min. Samples were subjected to a linear gradient of 2–25% solvent B over 120 minutes, followed by 25-40% B over 25 min, 40-90% B in 5min and finally 90% B over 10 min. The total runtime of the LC separation was180 minutes, including a 20 min column conditioning. The nanoLC system was coupled to an Orbitrap Lumos mass spectrometer via an EasySpray nano source (Thermo Fisher Scientific). The Orbitrap Lumos was operated in data-dependent acquisition mode using Xcalibur software. For MS2 acquisition, higher-energy collisional dissociation (HCD) MS/MS scans were recorded at a resolution of 60,000 following MS1 survey scans acquired at a resolution of 120,000. The MS1 ion target was set to 4 × 10⁵, and the MS2 target to 1 × 10⁵. Fragmentation was performed using a “Top Speed” acquisition strategy with a 3-second cycle time and a dynamic exclusion window of 45 seconds. The maximum ion injection time for MS2 scans was 105 ms, and the normalised HCD collision energy was set to 38. For MS3 acquisition, collision-induced dissociation (CID) MS2 scans were acquired at a resolution of 30,000, following an MS1 scan with the same parameters as above. The MS2 ion target was 5 × 10⁴, and multistage activation of the neutral loss (H₃PO₄) was enabled. The maximum injection time for MS2 scans was 60 ms, with a CID collision energy of 35. HCD MS3 scans were acquired at a resolution of 60,000, using synchronous precursor selection to include up to ten MS2 fragment ions. The MS3 ion target was 1 × 10⁵, with a maximum injection time of 105 ms and a collision energy of 65.

TMT-16plex: samples were resuspended in 0.1% TFA, 2% acetonitrile and loaded onto a 50 cm EasySpray PepMap column (75 μm inner diameter, 2 μm particle size; Thermo Fisher Scientific) fitted with an integrated electrospray emitter. Reverse-phase chromatography was performed using an RSLC nano U3000 system (Thermo Fisher Scientific) with a binary buffer system: solvent A consisted of 0.1% formic acid and 5% dimethyl sulfoxide (DMSO) in water, while solvent B contained 80% acetonitrile, 0.1% formic acid and 5% DMSO. Chromatographic separation was carried out at a flow rate of 250 nL/min. Samples were subjected to a linear gradient of 2–35% solvent B over 120 minutes, followed by 35-45% B over 25 min, 45-95% B in 5min and finally 95% B over 10 min. The total runtime of the LC separation was180 minutes, including a 20 min column conditioning. The nanoLC system was coupled to an Orbitrap Lumos mass spectrometer via an EasySpray nano source (Thermo Fisher Scientific).

The Orbitrap Lumos was operated in data-dependent acquisition mode using Xcalibur software. For MS2 acquisition, higher-energy collisional dissociation (HCD) MS/MS scans were recorded at a resolution of 50,000 following MS1 survey scans acquired at a resolution of 120,000. The MS1 ion target was set to 4 × 10⁵, and the MS2 target to 1 × 10⁵. Fragmentation was performed using a “Top Speed” acquisition strategy with a 3-second cycle time and a dynamic exclusion window of 45 seconds. The maximum ion injection time for MS2 scans was 120 ms, and the normalised HCD collision energy was set to 35. For MS3 acquisition, collision-induced dissociation (CID) MS2 scans were acquired at a resolution of 30,000, following an MS1 scan with the same parameters as above. The MS2 ion target was 5 × 10⁴, and multistage activation of the neutral loss (H₃PO₄) was enabled. The maximum injection time for MS2 scans was 60 ms, with a CID collision energy of 30. HCD MS3 scans were acquired at a resolution of 50,000, using synchronous precursor selection to include up to ten MS2 fragment ions. The MS3 ion target was 1 × 10⁵, with a maximum injection time of 120 ms and a collision energy of 55.

Raw mass spectrometry data were processed using MaxQuant ^81^ v1.6.2.10 (TMT-10plex) and v1.6.12.0 (TMT-16plex), with peptide identification carried out via the Andromeda ^82^ search engine. Spectra were searched against *Plasmodium falciparum* ^83^ and *Homo sapiens* ^84^ proteomes. TMT-based quantification was performed using the built-in ‘reporter ion MS2 or MS3’ algorithm in MaxQuant, with the reporter ion mass tolerance set to 0.003 Da. Carbamidomethylation of cysteine was set as a fixed modification, while variable modifications included methionine oxidation, protein N-terminal acetylation, and phosphorylation (S, T, Y). Enzyme specificity was set to trypsin, allowing up to two missed cleavages. The precursor mass tolerance was set to 20 ppm for the first search (used for mass recalibration) and 4.5 ppm for the main search. The ‘match between runs’ option was enabled (time window of 0.7 minutes) for fractionated samples. All datasets were filtered using posterior error probability (PEP) to achieve a 1% false discovery rate at the protein, peptide, and site levels.

Mass spectrometry datasets were imported into Perseus v1.5.0.9 ^85^ and filtered to exclude common contaminants as well as identifications derived from reverse decoy sequences. To generate a list of all quantified phosphorylation sites, reporter intensities were filtered to retain entries with at least one valid value. TMT reporter intensity values were log₂ transformed and normalized by median subtraction. Fold changes between conditions were then calculated for each experiment. Normalisation was performed in two steps. First, for each phosphosite, the mean log₂ intensity across all samples was subtracted from individual values to centre the data per site. Second, the median log₂ intensity of each sample was subtracted from all values in that sample to correct for global sample-level intensity differences ^86^. For each phosphosite quantified across all samples in a given experimental run, the mean log₂ intensity was calculated (i.e., the row mean). This mean was subtracted from each log₂ intensity value within that row. A median was then calculated for each column in the resulting matrix to determine a scaling factor for each sample. These median values were subtracted from the corresponding original log₂ intensity values within each column to obtain the final normalised intensities. The phosphoproteomic mass spectrometry data have been deposited to the ProteomeXchange Consortium via the PRIDE ^87^ partner repository with the dataset identifier PXD073843.

### Proteome analysis

Approximately 100 µg of the non-phosphorylated peptide flow-through was subjected to high-pH reversed-phase fractionation using the Pierce High pH Reversed-Phase Peptide Fractionation Kit (Thermo Fisher Scientific), following the manufacturer’s protocol for TMT-labeled samples with an additional 5% acetonitrile wash. Fractions were dried by vacuum centrifugation, stored at –80 °C, and subsequently analysed by LC-MS/MS on an Orbitrap Lumos mass spectrometer using data-dependent MS2 acquisition method as described above for TMT-10plex. Raw mass spectrometry data were processed using MaxQuant v1.6.2.10 ^81^ as described above with the following variable modifications: methionine oxidation, protein N-terminal acetylation, deamidation (NQ). Data were then imported into Perseus v1.5.0.9 ^85^ and filtered to exclude common contaminants as well as identifications derived from reverse decoy sequences and only identified by site. TMT reporter intensity values were log₂ transformed, normalized by median subtraction and filtered for one valid value to generate a list of quantified proteins.

### FIKK10.2-TurboID proxiome labelling

The protocol for TurboID-based proximity labelling and biotinylated peptide enrichment was previously described in detail by Davies *et al.* ^50^. TurboID-expressing parasites were cultured in biotin-free medium for two cycles prior to labelling. Parasites were tightly synchronised to a two-hour invasion window (>10% parasitaemia, 2% haematocrit in 300 mL total volume), and cultures were split into three experimental conditions, each performed in triplicate using red blood cells from independent donors (N = 3, 9 samples total).

Labelling was initiated by resuspending cells in medium containing 100 μM D-biotin (bioAPE); while not undergoing labelling, all cultures were maintained in biotin-free medium. Parasites were subjected to one of three total labelling conditions: continuous biotin labelling for 40 hours at 37 °C, or an 8-hour biotin pulse from 16–24 hpi at either 37 °C or 39 °C (heat stress). Outside these windows, biotin was removed by pelleting cultures and resuspending them in fresh biotin-free medium. At 40 hpi, samples were enriched by 60% Percoll gradient centrifugation and washed five times with PBS. Cells were lysed in ice-cold 8 M urea in 50 mM HEPES (pH 8.0), supplemented with cOmplete™ Mini, EDTA-free protease inhibitors (Roche). Lysates were sonicated (30% duty cycle, 3 × 30-second bursts on ice), clarified by centrifugation (13,000 × g, 30 minutes, 4 °C), snap-frozen in liquid nitrogen, and stored at −80 °C.

### Reduction, alkylation, and protein digest of biotinylated peptides

Parasite lysates were thawed and centrifuged to pellet insoluble material (15 minutes, 4 °C, 21,000 × g). Protein content in the supernatant was quantified using the BCA Protein Assay Kit (Pierce). An equal amount of parasite material (4 mg per sample) was taken, and protein concentrations were equalised using 8 M urea in 50 mM HEPES. Oxidised cysteine residues were reduced with 5 mM dithiothreitol (DTT) at room temperature for 60 minutes, followed by alkylation of the reduced cysteines with 10 mM iodoacetamide in the dark at room temperature for 30 minutes. For protein digestion, samples were diluted 4-fold with 50 mM HEPES to reduce the urea concentration to below 2 M. Mass spectrometry-grade trypsin was added at a 1:50 enzyme-to-protein ratio and incubated at 37 °C overnight (16 hours). To terminate digestion, samples were cooled on ice for 10 minutes and acidified with TFA to a final concentration of ∼0.4% (v/v), incubating on ice for a further 10 minutes. Samples were centrifuged at 21,000 × g for 15 minutes at 4 °C to remove insoluble material. These peptides were then Sep-Pak desalted and vacuum-dried as previously described.

### Charging protein G agarose beads with anti-biotin antibodies

For each sample, 60 µL of a 50% protein-G agarose bead slurry (Pierce #20398) was used and washed three times with 5–10 bead volumes of BioSITe buffer ^88^ comprising 50 mM Tris, 150 mM NaCl, and 0.5% Triton X-100 (pH 7.2–7.5 at 4 °C). Between washes, the beads were pelleted by centrifugation at 1500 × g for 2 minutes at 4 °C. Anti-biotin antibodies (a mixture of 30 µg each of Bethyl Laboratories #150-109A and Abcam #Ab53494) was added per 60 µL of bead slurry, brought up to 2 mL with BioSITe buffer, and incubated overnight at 4 °C with rotation. The beads were then washed three times with BioSITe buffer, adjusted to approximately a 50% slurry using the same buffer, and aliquoted into 60 µL portions for each sample.

### Biotinylated peptide immunoprecipitation

Dried peptide samples were reconstituted in 1.5 mL of BioSITe buffer with vortexing to facilitate dissolution. The pH was adjusted to between 7 and 7.5 on ice using 1–5 µL of 10 M NaOH. Samples were then centrifuged at 21,000 × g for 10 minutes at 4 °C to remove any insoluble material. Peptide concentrations were determined using the peptide BCA assay kit (Pierce, #23275). Equal peptide amounts from each sample were added to the anti-biotin antibody-conjugated bead slurry (∼60 µL per sample) and incubated with rotation for 2 hours at 4 °C. Following incubation, beads were pelleted by centrifugation (1500 × g, 2 minutes, 4 °C) and sequentially washed with 3 × 0.5 mL BioSITe buffer, 1 × 0.5 mL of 50 mM Tris, and 3 × 1 mL of mass spectrometry-grade water. Biotinylated peptides were eluted by treating the beads with 4 × 50 µL of 0.2% TFA, and eluates were stored at –80 °C.

### LC-MS/MS (TurboID biotinylated peptide pulldown) and data analysis

Samples were loaded onto Evotips (according to the manufacturer’s instructions). Following a wash with aqueous acidic buffer (0.1% formic acid in water), samples were loaded onto an Evosep One system coupled to an Orbitrap Fusion Lumos (Thermo Fisher Scientific). The Evosep One was fitted with a 15 cm column (PepSep) and a predefined gradient for a 44 min method was employed. The Orbitrap Lumos was operated in data-dependent mode (1 s cycle time), acquiring IT HCD MS/MS scans in rapid mode after an OT MS1 survey scan (*R* = 60,000). The MS1 target was 4E5 ions, whereas the MS2 target was 1E4 ions. The maximum ion injection time utilised for MS2 scans was 300 ms, the HCD normalised collision energy was set at 32, and the dynamic exclusion was set at 15 s.

Acquired raw files were processed with MaxQuant v1.6.2.10 ^81^. Peptides were identified from the MS/MS spectra searched against *P. falciparum* ^83^ and Homo sapiens ^84^ proteomes using Andromeda ^82^ search engine. Biotin (K), Oxidation (M), Acetyl (Protein N-term) and Phospho (STY) were selected as variable modifications whereas Carbamidomethyl (C) was selected as a fixed modification. The enzyme specificity was set to trypsin with a maximum of three missed cleavages. Minimal peptide length was set at 6 amino acids. Biotinylated peptide search in Max Quant was enabled by defining a biotin adduct (+226.0776) on lysine residues as well as its three diagnostic ions: fragmented biotin (m/z 227.0849), immonium ion harboring biotin with a loss of NH3 (m/z 310.1584), and an immonium ion harboring biotin (m/z 327.1849). The precursor mass tolerance was set to 20 ppm for the first search (used for mass re-calibration) and to 4.5 ppm for the main search. The datasets were filtered on posterior error probability (PEP) to achieve a 1% false discovery rate on protein, peptide and site level. Other parameters were used as pre-set in the software.

MaxQuant output files were processed with Perseus, v1.5.0.9 ^85^, the data were filtered to exclude common contaminants as well as identifications derived from reverse decoy sequences. Peptide intensities were log₂-transformed, and peptides were filtered to retain those detected in two out of three replicates within at least one condition. Valid peptides were assigned for each group as being present in at least 2 of 3 replicate conditions. Valid proteins were defined for each condition as having at least 2 valid peptides. Valid peptides and proteins from each condition were used to generate area-proportional Venn diagrams using the BioVenn tool ^89^. The TurboID mass spectrometry data have been deposited to the ProteomeXchange Consortium via the PRIDE ^87^ partner repository with the dataset identifier PXD073890.

### Statistics and reproducibility

Data were analysed using GraphPad Prism v10 and details of the analyses are provided in each figure legend. Where specified, data were log transformed prior to statistical testing. Percentage data constrained between 0 and 100% were first converted to proportions and then Logit transformed. Fold change data were Log10 transformed as indicated. Multiple comparisons were corrected for false discovery rate (FDR) using the method of Benjamini Krieger and Yekutieli. Where FDR correction was applied, adjusted P values (Q values) are reported. Statistical significance was defined as follows: ns not significant, *P < 0.05, **P < 0.01, ***P < 0.001, ****P < 0.0001.

## List of Supplementary Figures and Tables

- Figure 1 - Supplement 1. Heat stress applied between 16-24 hours post-invasion does not accelerate *P. falciparum* asexual development.
- Figure 1 - Supplement 2. Flow cytometry gating strategy for detection of surface-trafficked VAR2CSA on iRBCs.
- Figure 1 - Supplement 3. Temperature probe within cell culture flasks accurately measures heat stress conditions applied to cell cultures.
- Figure 2 - Supplement 1. Sorbitol lysis of infected red blood cells is reduced by the PSAC inhibitor furosemide regardless of temperature treatment.
- Figure 2 - Supplement 2. *P. falciparum* parasites are not more susceptible to heat stress with reduced nutrient availability.
- Figure 3 - Supplement 1. The host and parasite proteomes are not substantially altered following febrile heat stress.
- Figure 3 - Supplement 2. Repeated heat stress phosphoproteome with FIKK knockout (KO) strains identifies FIKK10.2 as the major contributor of heat stress dependent phosphorylation of host-cell exported parasite proteins.
- Figure 3 - Supplement 3. Generation of HSP70x-3xHA (PF3D7_0831700) conditional knockout *P. falciparum* strain.
- Figure 3 - Supplement 4. HSP70x-3xHA (PF3D7_0831700) does not strongly co-localise with KAHRP or SBP1.
- Figure 3 - Supplement 5. Generation and validation of PF3D7_0702500-3xHA conditional knockout *P. falciparum* strain.
- Figure 3 - Supplement 6. PF3D7_0831700-3xHA co-localises with SBP1.
- Figure 3 - Supplement 7. HSP70x conditional knockout parasites have normal growth at 37 °C and elevated temperatures.
- Figure 3 - Supplement 8. The exported protein HSP70x is not required for the trafficking of PfEMP1 onto the infected cell’s surface under normal or febrile temperatures.
- Figure 3 - Supplement 9. HSP70x is not required for functional PSAC activity at 24 hpi under normal or elevated temperatures.
- Figure 3 - Supplement 10. PF3D7_0702500 conditional knockout parasites have normal growth at 37 °C and elevated temperatures.
- Figure 3 - Supplement 11. The exported protein PF3D7_0702500 is not required for the trafficking of PfEMP1 onto the infected cell’s surface under normal or febrile temperatures.
- Figure 3 - Supplement 12. PF3D7_0702500 is not required for functional PSAC activity at 24 hpi under normal or elevated temperatures.
- Figure 4 - Supplement 1. Generation and validation of FIKK10.2-TurboID *P. falciparum* transgenic strain.
- Figure 4 - Supplement 2. Predicted Maurer’s cleft protein topology from enriched phosphopeptides and biotinylated peptides.
- Figure 5 - Supplement 1. Generation and validation of VAR2CSA-NanoLuc-3xHA *P. falciparum* transgenic strain.
- Figure 6 - Supplement 1. Generation and validation of constitutively expressed NanoLuc reporter strains.
- Supplementary Table 1. Heat stress phosphoproteome and proteome data from the 10-plex TMT from NF54::DiCre (wild type). Includes *P. falciparum* and *H. sapiens* peptides and proteins.
- Supplementary Table 2. *P. falciparum* phosphosites that are more phosphorylated following heat stress.
- Supplementary Table 3. Heat stress phosphoproteome and proteome data from the 16-plex TMT with FIKK conditional knockouts. Includes *P. falciparum* and *H. sapiens* peptides and proteins.
- Supplementary Table 4. FIKK10.2-TurboID proximal biotinylated proteins identified under various conditions, including the presence or absence of biotin and heat stress.
- Supplementary Table 5. FIKK10.2-TurboID proximal biotinylated proteins identified in the presence of biotin and in the absence of heat stress
- Supplementary Table 6. Primer sequences and binding sites used for validation of transgenic *P. falciparum* strains.
- Supplementary Table 7. Composition of reduced nutrient media

## Notes

### Competing Interest Statement

The authors have declared no competing interest.

### Summary of Updates

The manuscript has been updated following reviewers comments and now includes additional data to support the sorbitol sensitivity experiments (Figure 2 - Supplement 1) and to assess the role of nutrient uptake in P. falciparum growth following febrile temperatures (Figure 2 - Supplement 2). The updated manuscript has also undergone reformatting and includes updated microscopy images (Figure 3 - Supplement 4 and Figure 3 - Supplement 6).

