## Supplementary Figures for "Physiological febrile heat stress increases cytoadhesion through increased protein trafficking of *Plasmodium falciparum* surface proteins into the red blood cell"

**Figure 1 -Supplement 1.** Heat stress applied between 16-24 hours post-invasion does not accelerate *P. falciparum asexual* development.

**Figure 1 - Supplement 2.** Flow cytometry gating strategy for detection of surface-trafficked VAR2CSA on iRBCs.

**Figure 1 - Supplement 3.** Temperature probe within cell culture flasks accurately measures heat stress conditions applied to cell cultures.

**Figure 2 - Supplement 1.** Sorbitol lysis of infected red blood cells is reduced by the PSAC inhibitor furosemide regardless of temperature treatment.

**Figure 2 - Supplement 2***. P. falciparum parasites* are not more susceptible to heat stress with reduced nutrient availability.

**Figure 3 - Supplement 1.** The host and parasite proteomes are not substantially altered following febrile heat stress.

**Figure 3 - Supplement 2.** Repeated heat stress phosphoproteome with FIKK knockout (KO) strains identifies FIKK10.2 as the major contributor of heat stress dependent phosphorylation of host-cell exported parasite proteins.

**Figure 3 - Supplement 3.** Generation of HSP70x-3xHA (PF3D7_0831700) conditional knockout *P. falciparum* strain.

**Figure 3 - Supplement 4.** HSP70x-3xHA (PF3D7_0831700) does not strongly co-localise with KAHRP or SBP1.

**Figure 3 - Supplement 5.** Generation and validation of PF3D7_0702500-3xHA conditional knockout *P. falciparum* strain.

**Figure 3 - Supplement 6.** PF3D7_0831700-3xHA co-localises with SBP1.

**Figure 3 - Supplement 7.** HSP70x conditional knockout parasites have normal growth at 37 °C and elevated temperatures.

**Figure 3 - Supplement 8.** The exported protein HSP70x is not required for the trafficking of PfEMP1 onto the infected cell's surface under normal or febrile temperatures.

**Figure 3 - Supplement 9.** HSP70x is not required for functional PSAC activity at 24 hpi under normal or elevated temperatures.

**Figure 3 - Supplement 10.** PF3D7_0702500 conditional knockout parasites have normal growth at 37 °C and elevated temperatures.

**Figure 3 - Supplement 11.** The exported protein PF3D7_0702500 is not required for the trafficking of PfEMP1 onto the infected cell's surface under normal or febrile temperatures.

**Figure 3 - Supplement 12**. PF3D7_0702500 is not required for functional PSAC activity at 24 hpi under normal or elevated temperatures.

**Figure 4 - Supplement 1.** Generation and validation of FIKK10.2-TurboID *P. falciparum* transgenic strain.

**Figure 4 - Supplement 2.** Predicted Maurer's cleft protein topology from enriched phosphopeptides and biotinylated peptides.

**Figure 5 - Supplement 1.** Generation and validation of VAR2CSA-NanoLuc-3xHA *P. falciparum* transgenic strain.

**Figure 6 - Supplement 1.** Generation and validation of constitutively expressed NanoLuc reporter strains.

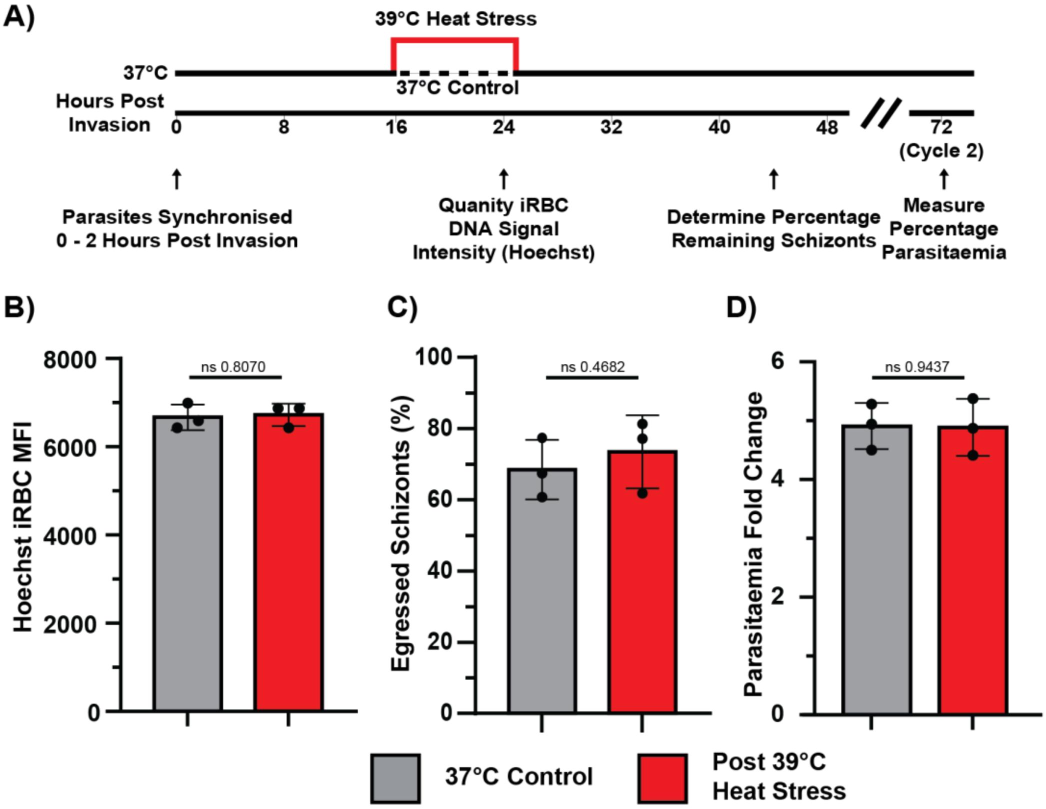


**Figure 1 - Supplement 1. Heat stress applied between 16-24 hours post-invasion does not accelerate *P. falciparum* asexual development.** (A) Schematic overview of the experimental design. (B) Median fluorescent intensity (MFI) of Hoechst-stained infected red blood cells (iRBCs) at 24 hours post-infection (hpi) was not significantly altered by heat stress. (C) High DNA content iRBCs, identified as schizonts, showed no significant difference in proportion remaining at 44 hpi following heat stress. (D) Fold increase in parasitaemia in the subsequent asexual cycle was comparable between heat-stressed samples and the 37 °C control. Data represent N = 3 biological replicates. Error bars indicate ±1 standard deviation (SD). Statistical comparisons were made using unpaired t-tests with Welch's correction. For panels (C) and (D), data were log-transformed prior to analysis.

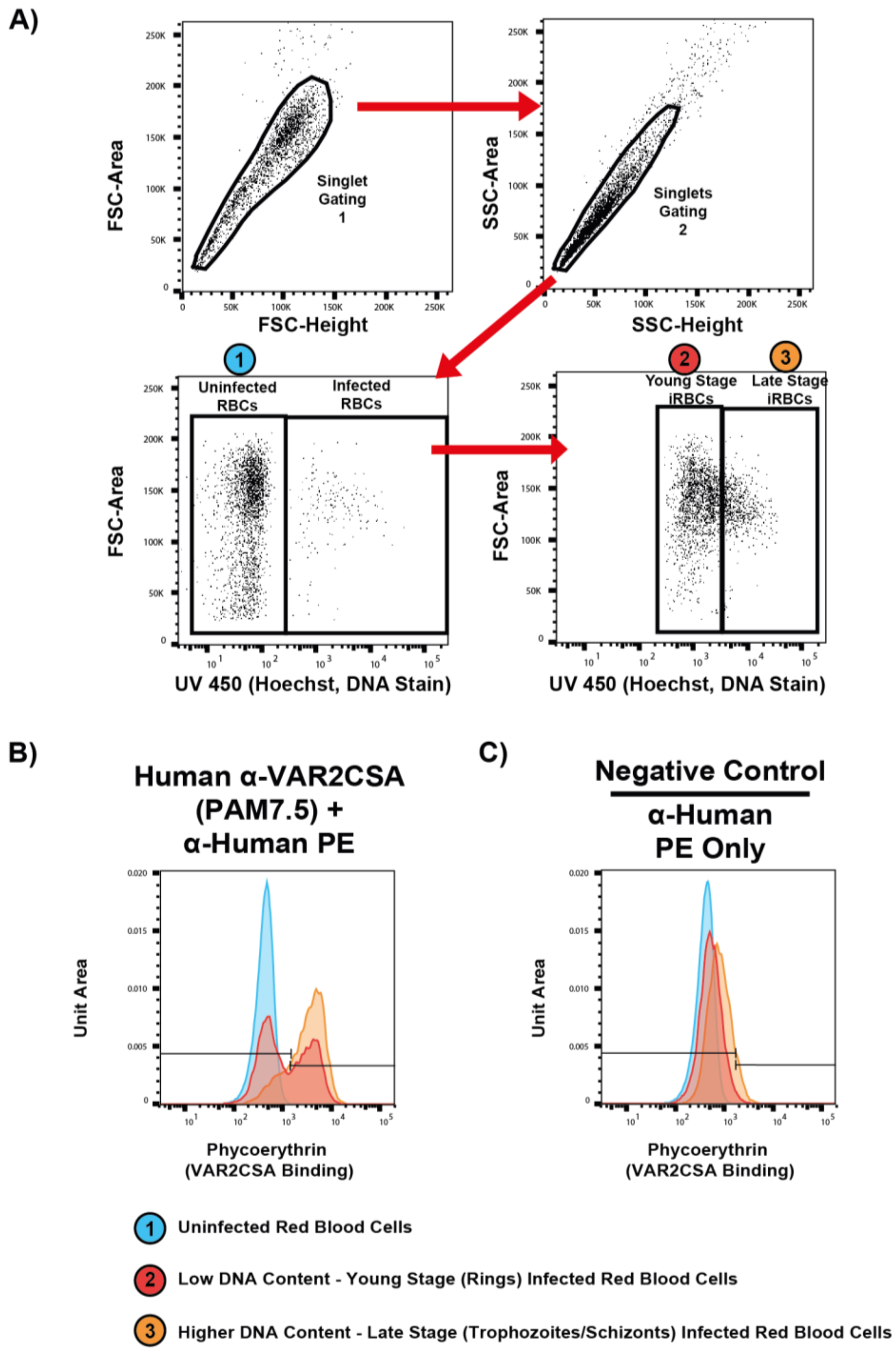


**Figure 1 - Supplement 2. Flow cytometry gating strategy for detection of surface-trafficked VAR2CSA on iRBCs.** (A) Example flow cytometry gating strategy for NF54::DiCre parasites. Singlets were gated, followed by DNA staining to distinguish uninfected and infected red blood cells. Infected red blood cells were further gated into young (lower DNA content) and late stage (higher DNA content) iRBCs. (B) iRBCs show increased signal following incubation with human anti-VAR2CSA antibody (PAM7.5) and phycoerythrin-conjugated secondary antibody. (C) A negative control, in the absence of primary antibody but with the secondary fluorophore, shows reduced signal on iRBCs. The iRBC gate including all stages was used to determine the percentage of cells positive for PfEMP1 surface labelling.

A)

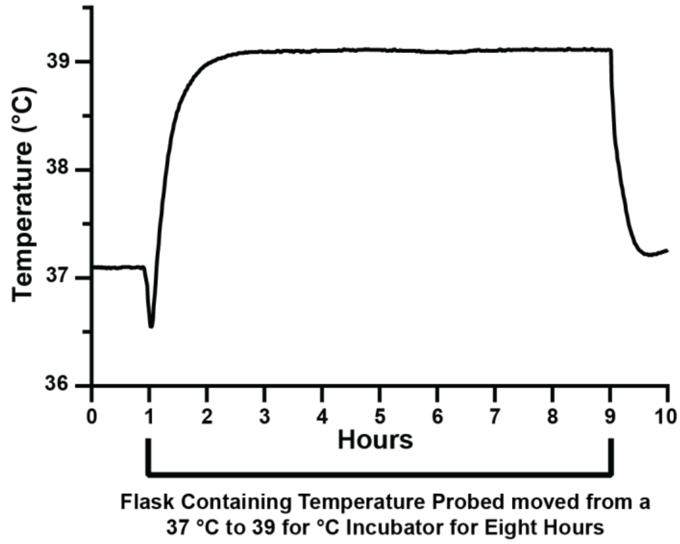


**Figure 1 - Supplement 3. Temperature probe within cell culture flasks accurately measures heat stress conditions applied to cell cultures.** (A) Thermal profile of the *in vitro* heat stress protocol. A 10 mL water-filled flask was transferred from a 37 °C to a 39 °C incubator, and temperature was recorded every 30 seconds using a probe (±0.2 °C) placed in the liquid. The flask remained at 39 °C for 8 hours before being returned to 37 °C.

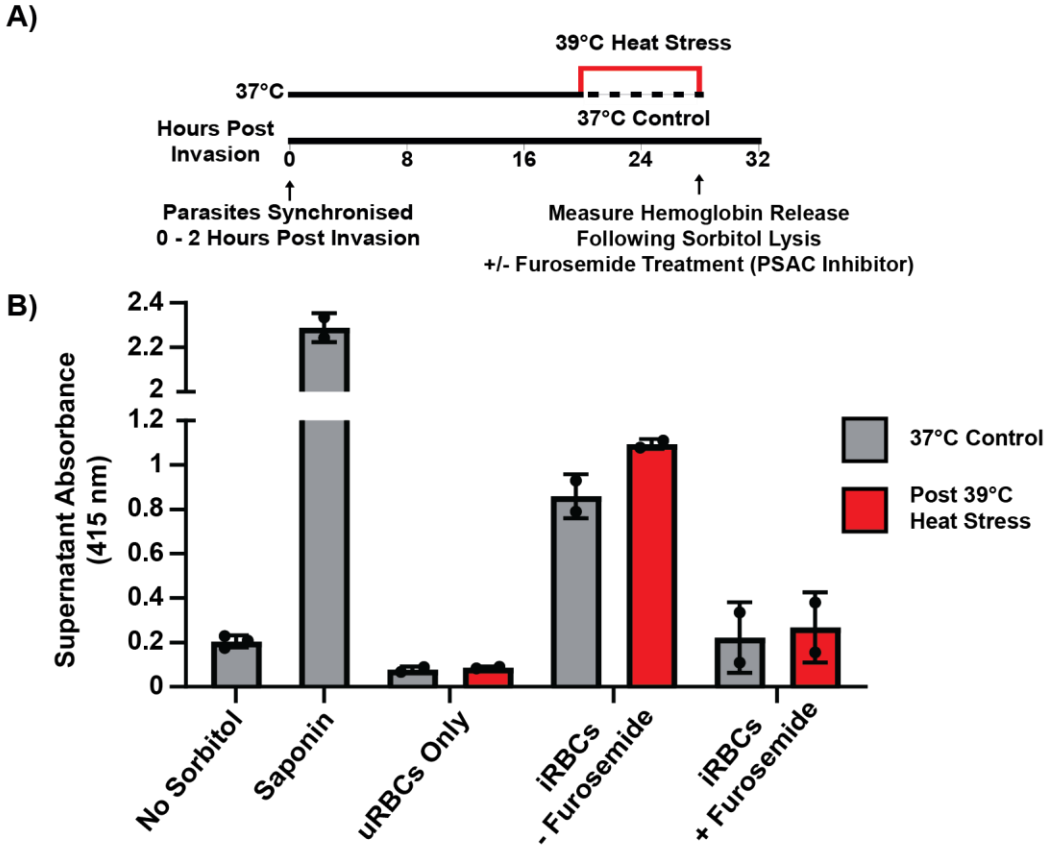


**Figure 2 - Supplement 1. Sorbitol lysis of infected red blood cells is reduced by the PSAC inhibitor furosemide regardless of temperature treatment.** (A) Schematic overview of the experimental workflow. NF54 *P. falciparum* infected red blood cells (iRBCs) were synchronised to a 2-hour window (>10% parasitaemia). At 20 hours post infection, cultures were either maintained at 37 °C or exposed to heat stress at 39 °C for 8 hours. At 28 hours post infection, all cells were washed with room-temperature phosphate-buffered saline and a subset of samples was pre-incubated with 200 µM furosemide for 10 minutes at 37 °C. Samples were then resuspended in sorbitol or sorbitol plus furosemide and incubated for 10 minutes at 37 °C. Following incubation, cells were centrifuged and supernatant absorbance at 415 nm was measured as an indicator of hemoglobin release and therefore cell lysis. Uninfected red blood cells (uRBCs) were processed in parallel. Sorbitol uptake is proposed to occur through the plasmodial surface anion channel (PSAC), which is inhibited by furosemide. (B) Supernatant absorbance at 415 nm of sorbitol-treated cultures following heat stress compared with the non heat-stressed 37 °C control. Negative and positive supernatant controls were generated from iRBCs treated without sorbitol and from iRBCs lysed with saponin (supernatant diluted 1:5) respectively. Only iRBCs showed increased supernatant absorbance after sorbitol treatment, and this response was blocked by furosemide. uRBCs showed no increase in absorbance, indicating that sorbitol-mediated lysis requires parasite infection. Data represent at least two biological replicates. Error bars indicate ±1 standard deviation (SD).

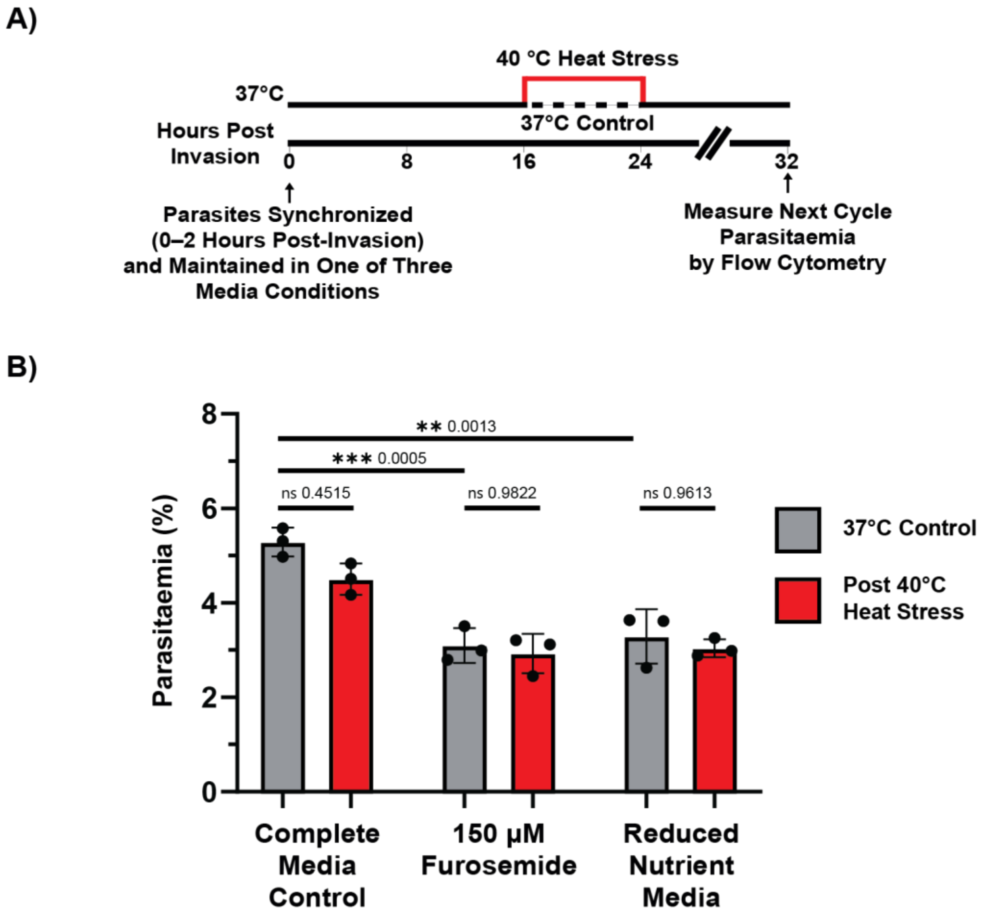


**Figure 2 - Supplement 2. *P. falciparum* parasites are not more susceptible to heat stress with reduced nutrient availability.** A) NF54 DiCre parasites were synchronised to a two-hour invasion window and adjusted to 1% parasitaemia before culture with complete media (supplemented RPMI-1640), complete media containing 150 µM furosemide, or reduced nutrient media (composition detailed in Supplementary Table 7). Between 16-24 hpi iRBCs were heat stressed at 40 °C or maintained at 37 °C, in the following cycle (72 hpi) parasitaemia was determined. B) Culturing with 150 µM furosemide, an inhibitor that blocks nutrient uptake, or with reduced nutrient media resulted in a significant reduction in replication compared to the complete media control. However, these reductions were not further decreased during heat stress. Data represent N = 3 biological replicates. Error bars indicate ±1 standard deviation (SD). Both the normal media control and the Reduced nutrient media were treated with a DMSO vehicle control. Statistical significance was assessed using two-way ANOVA of log-transformed data, followed by FDR correction for multiple comparisons by Šidák's correction.

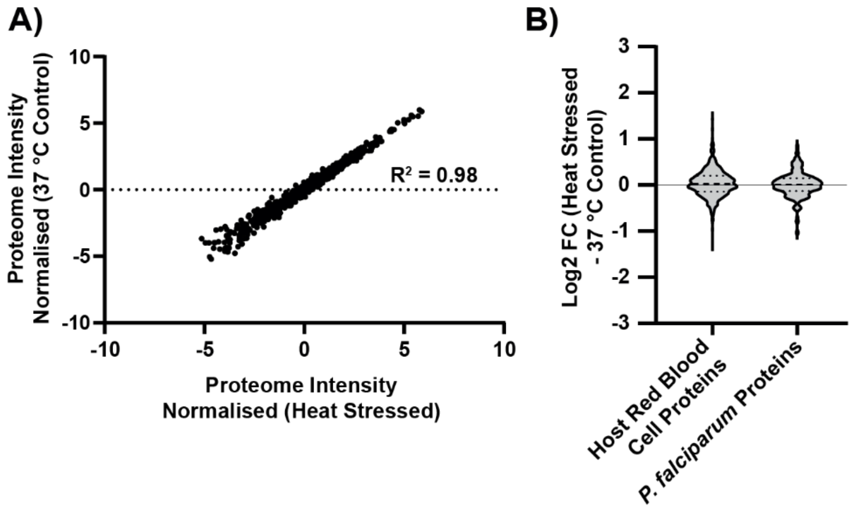


**Figure 3 - Supplement 1. The host and parasite proteomes are not substantially altered following febrile heat stress.** (A) Normalised protein abundance of human (n = 212) and Plasmodium falciparum (n = 624) proteins from heat-stressed and 37 °C control infected red blood cells (iRBCs), totalling 836 proteins. Pearson correlation analysis between conditions is shown ($R^2 = 0.98$). (B) Distribution of $\log_2$ fold changes in protein abundance for human (n = 212) and *P. falciparum* (n = 624) proteins comparing heat-stressed versus 37 °C control iRBCs. Data are from two biological replicates. Violin plots display the median and interquartile range (25th and 75th percentiles, indicated by dotted lines).

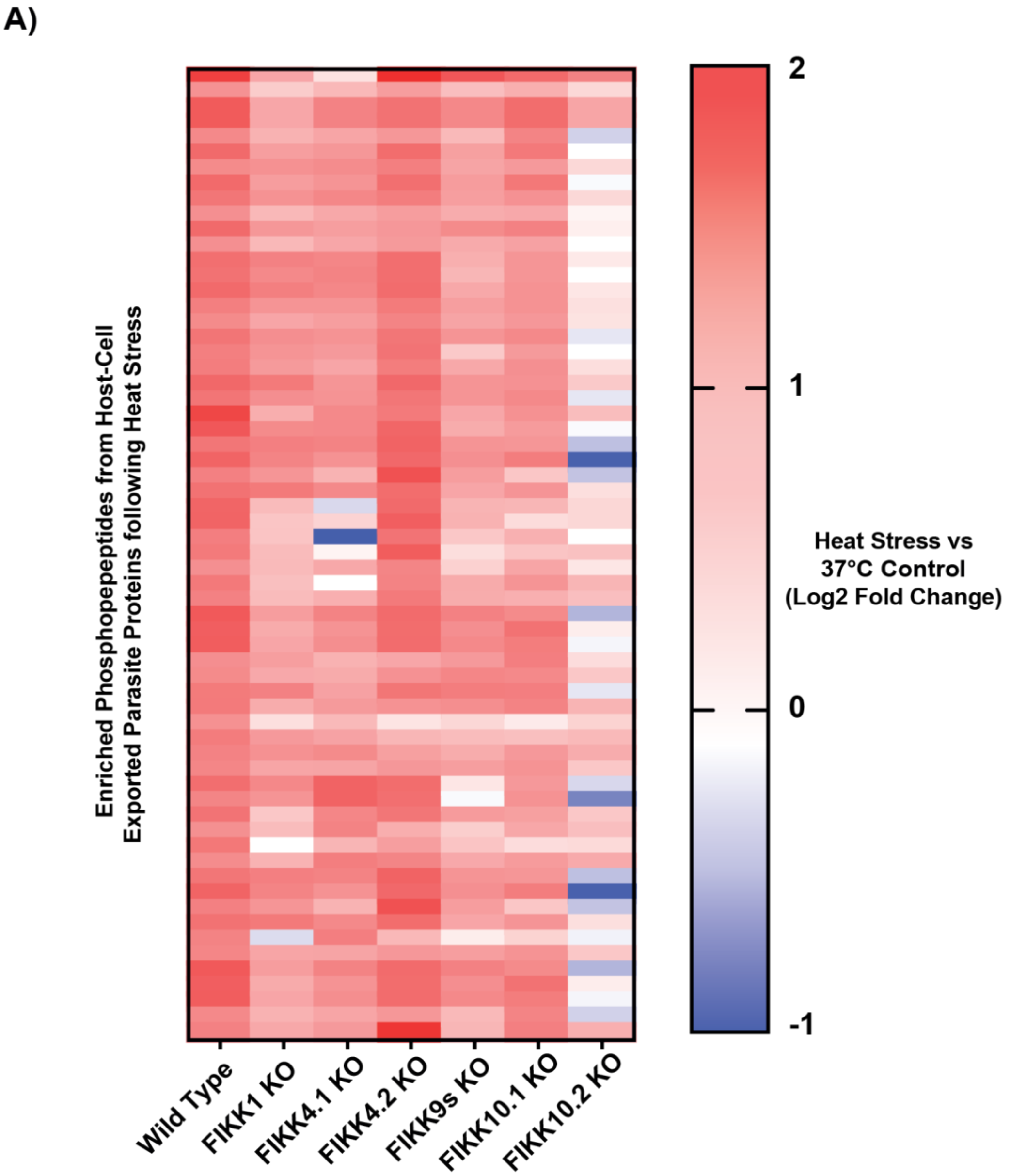


**Figure 3 - Supplement 2. Repeated heat stress phosphoproteome with FIKK knockout (KO) strains identifies FIKK10.2 as the major contributor of heat stress dependent phosphorylation of host-cell exported parasite proteins.** (A) Heat map of 64 exported phosphopeptides that were significantly more abundant following heat stress in wild-type–like NF54 DiCre parasites ($\log_2$ fold change > 1; heat stress vs 37 °C control). Fold changes for these phosphopeptides are shown across several exported FIKK kinase conditional knockouts (KO). The FIKK9s strain represents the conditional deletion of seven FIKK kinases (FIKK9.1 to FIKK9.7). In the absence of FIKK10.2, 52 of the 64 phosphopeptides that were significantly more phosphorylated under heat stress in the wild-type strain no longer show increased phosphorylation. N = 1 for each condition (temperature and strain).

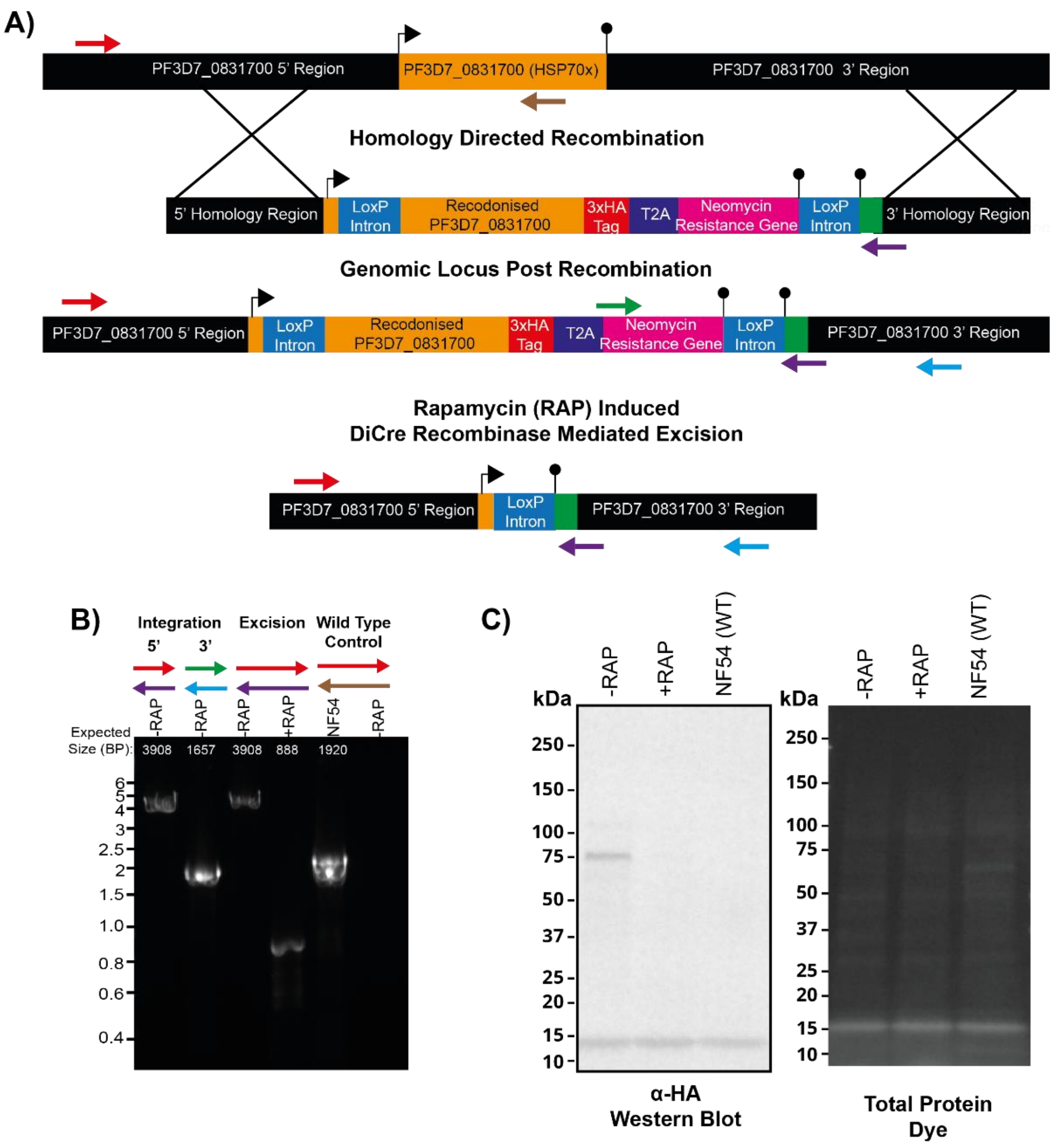


**Figure 3 - Supplement 3. Generation of HSP70x-3xHA (PF3D7_0831700) conditional knockout *P. falciparum* strain.** A) Generation of a conditional knockout strain of HA-tagged HSP70x using CRISPR/Cas9-mediated excision and homologous repair. Upon rapamycin (+RAP) treatment, Cre recombinase excises DNA flanked by loxP sites to remove HSP70x. Primer binding sites indicated by the red and blue arrows are outside the 5' and 3' repair template homology regions. B) PCR confirmed 5′ and 3′ integration, reduction in gene-specific amplification following deletion, and absence of the original wild-type (WT) allele. Predicted amplicon sizes are indicated in white. DNA size marker values (kbp) are indicated to the left of the gel. C) Western blot using anti-HA antibody of Percoll-enriched HSP70x-3xHA schizonts previously treated with DMSO (−RAP) or rapamycin (+RAP), compared to WT control (NF54). Total protein staining was used as a loading control. The predicted size of HSP70x-3xHA is 80.4 kDa.

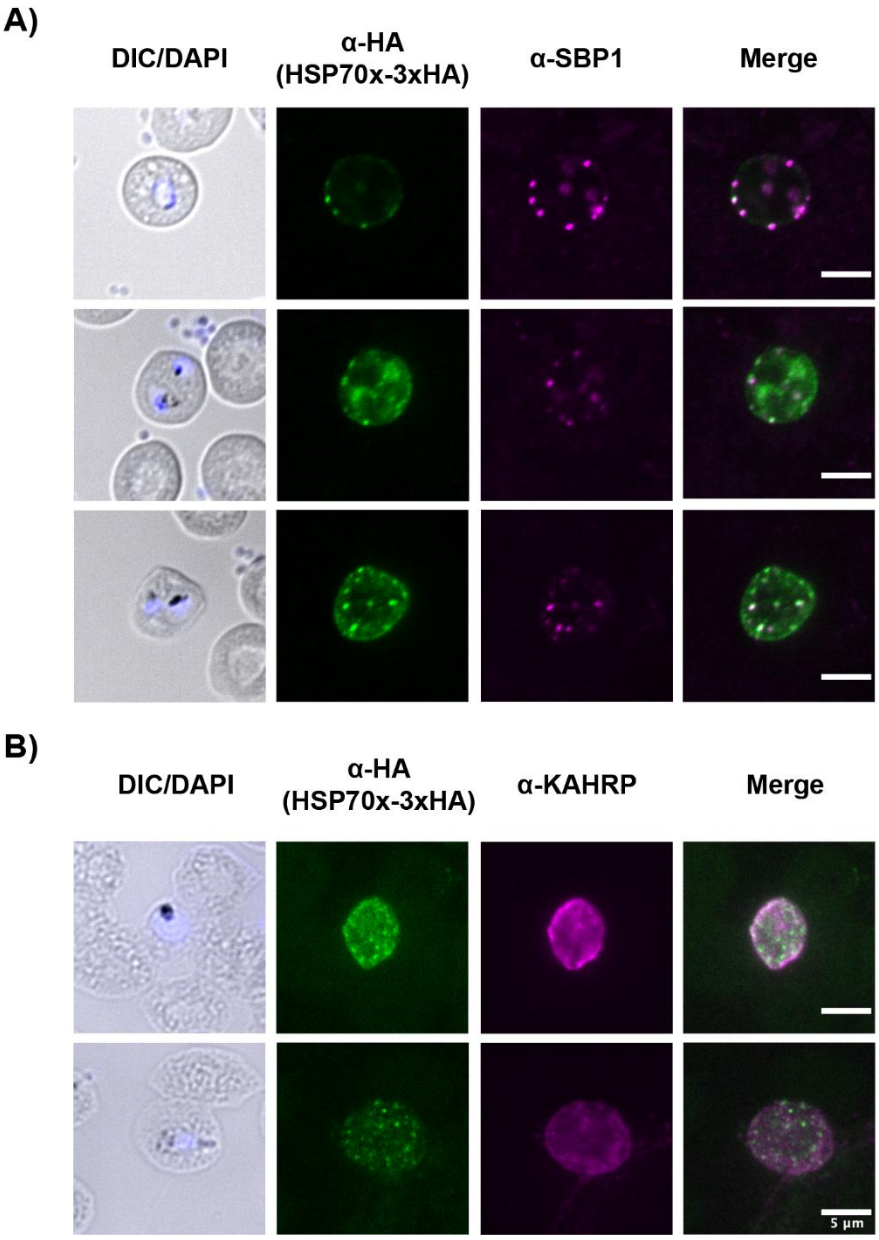


**Figure 3 - Supplement 4. HSP70x-3xHA (PF3D7_0831700) does not strongly co-localise with KAHRP or SBP1.** Representative immunofluorescence microscopy images showing HSP70x-3xHA (anti-HA), parasite DNA (DAPI) merged with differential interference contrast (DIC), and either KAHRP (A) or SBP1 (B). Neither KAHRP (a red blood cell cytoskeleton-associated protein) nor SBP1 (a Maurer's cleft-resident protein) co-localised with HA-tagged HSP70x. Scale bar = 5 µm.

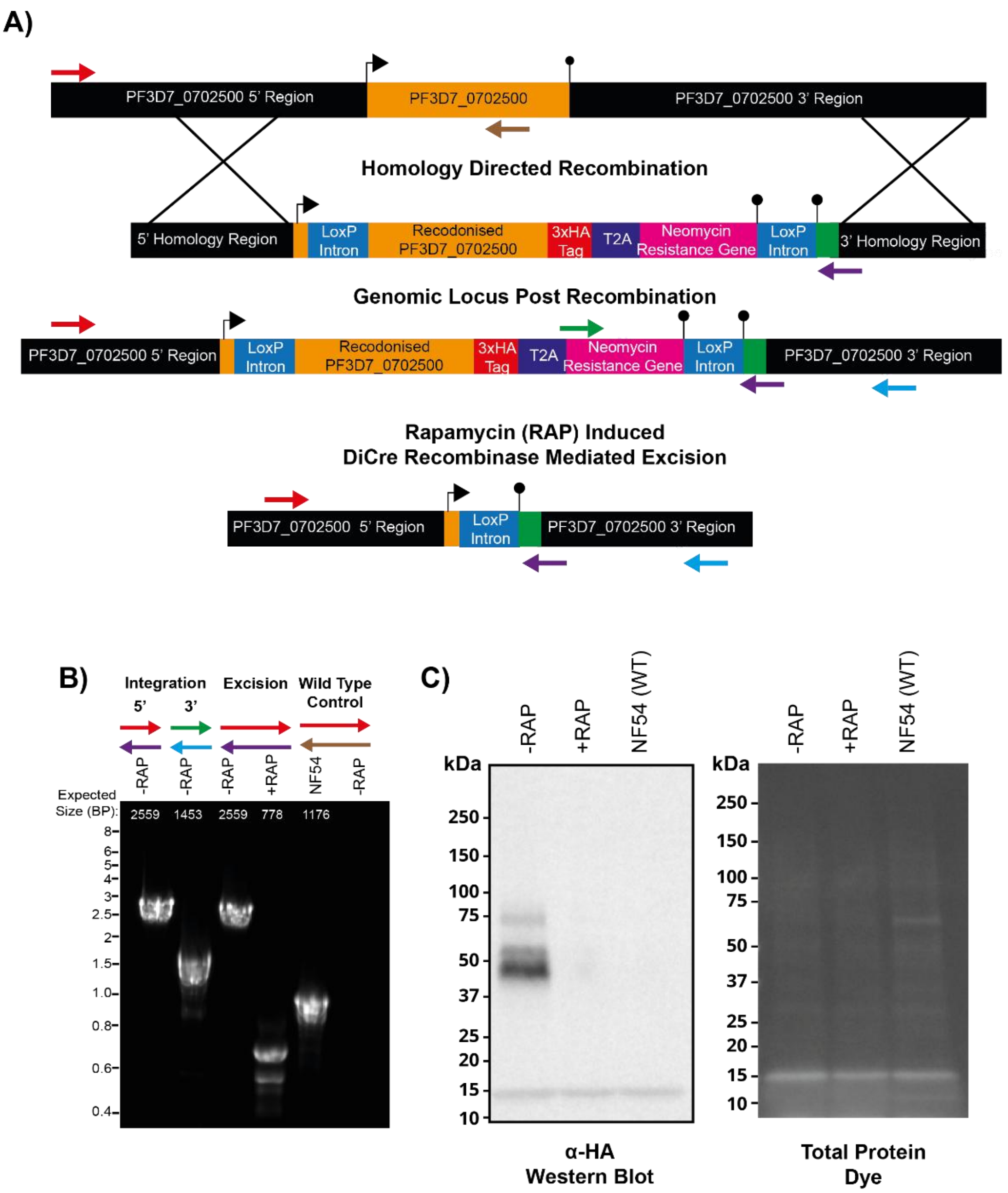


**Figure 3 - Supplement 5. Generation and validation of PF3D7_0702500-3xHA conditional knockout *P. falciparum* strain.** A) Generation of a conditional knockout strain of HA-tagged PF3D7_0702500 using CRISPR/Cas9-mediated excision and homologous repair. Upon rapamycin (+RAP) treatment, Cre recombinase excises DNA flanked by loxP sites to remove PF3D7_0702500. Primer binding sites indicated by the red and blue arrows are outside the 5' and 3' repair template homology regions. B) PCR confirmed 5′ and 3′ integration, reduction in gene-specific amplification following deletion, and absence of the original wild-type (WT) allele. Predicted amplicon sizes are indicated in white. DNA size marker values (kbp) are indicated to the left of the gel. C) Western blot using anti-HA antibody of Percoll-enriched PF3D7_0702500-3xHA schizonts previously treated with DMSO (−RAP) or rapamycin (+RAP), compared to WT control (NF54). Total protein staining was used as a loading control. The predicted size of PF3D7_0702500-3xHA is 33.0 kDa. The observed discrepancies in apparent molecular weight may result from extensive post-translational modifications and are consistent with findings previously reported by Heiber *et al.* [54]. Additionally, the faint larger band likely represents a population in which the T2A skip peptide failed to cleave efficiently, resulting in a fusion with the neomycin resistance cassette. This fusion protein is predicted to be 62.1 kDa (an additional 29.1 kDa).

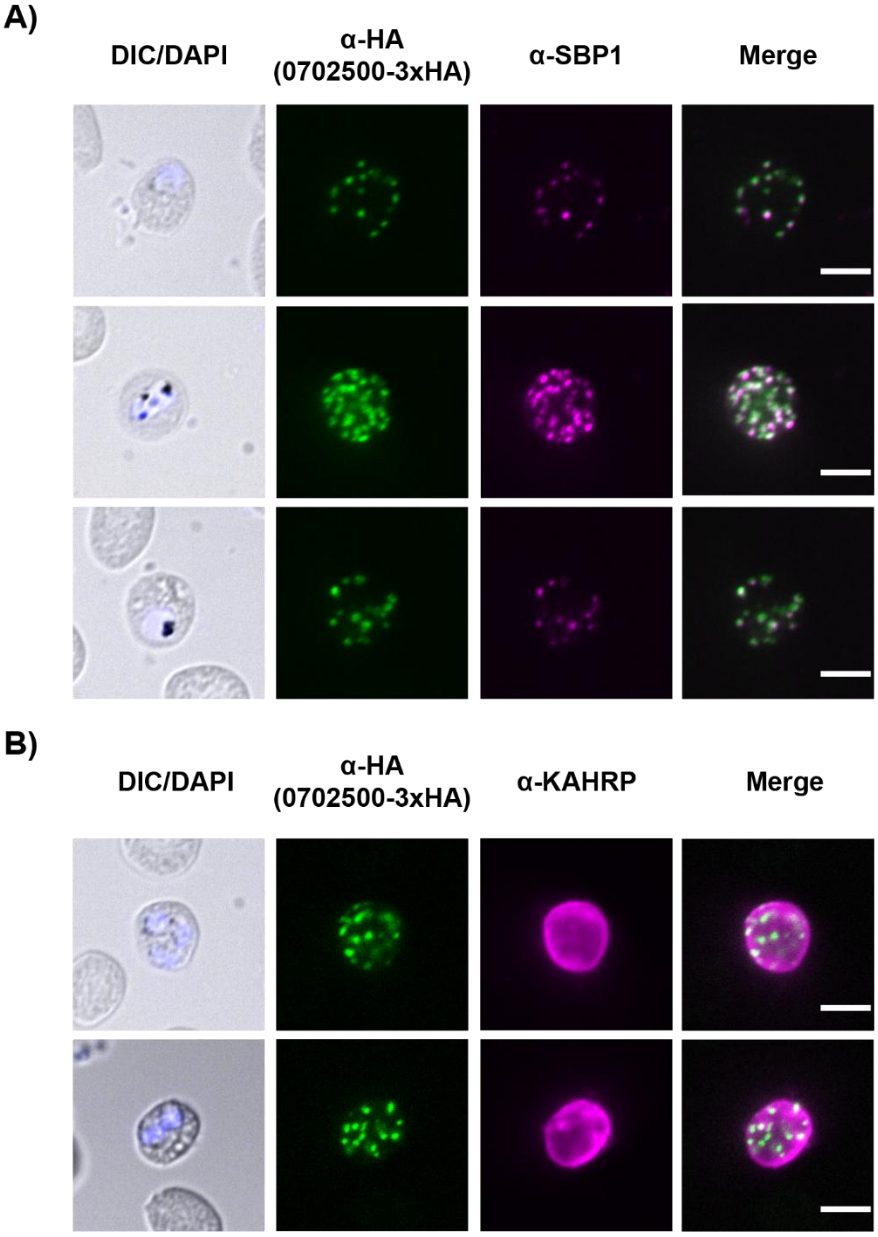


**Figure 3 - Supplement 6. PF3D7_0702500-3xHA co-localises with SBP1.** Representative immunofluorescence microscopy images showing PF3D7_0702500-3xHA (anti-HA), parasite DNA (DAPI) merged with differential interference contrast (DIC), and either KAHRP (A) or SBP1 (B). PF3D7_0702500-3xHA does not co-localise with KAHRP (a red blood cell cytoskeleton-associated protein) but does co-localise with SBP1 (a Maurer's cleft-resident protein). Scale bar = 5 µm.

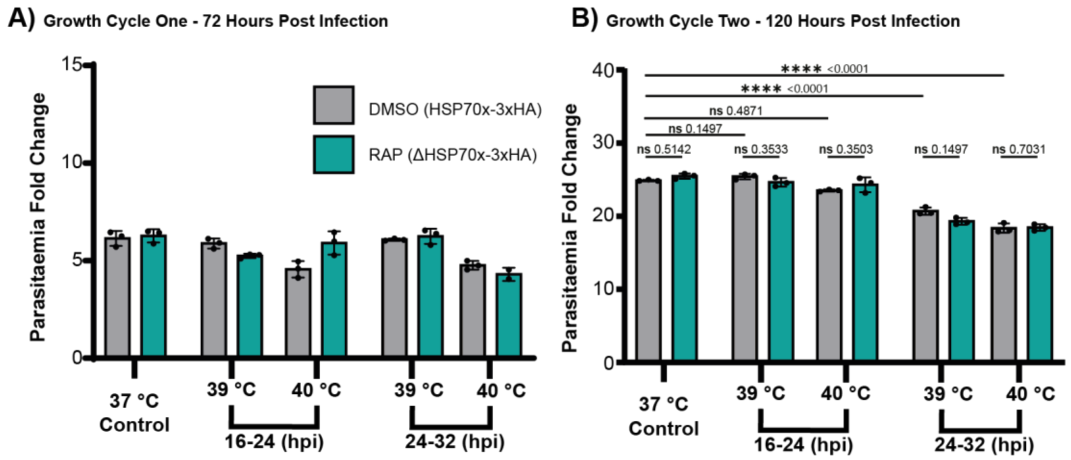


**Figure 3 - Supplement 7. HSP70x conditional knockout parasites have normal growth at 37 °C and elevated temperatures.** Parasites were either DMSO or rapamycin (RAP) treated (four hours) to conditionally knockout HSP70x (PF3D7_0831700), parasites were later synchronised to a two-hour invasion window. Parasites were heat stressed at 39 °C or 40 °C between 16-24, 24-32 hpi or kept at 37 °C for the full duration. 40 °C heat stress between 24-32 hpi significantly reduced growth as shown in the fold change in parasitaemia in the first (A) and second cycle (B) following heat stress. However, the absence of HSP70x (RAP treated) did not have a significant impact on the growth of the parasites following any of the heat stress conditions tested compared to the wildtype resembling (DMSO treated) strain. Data represent N=3 biological replicates, with error bars showing ±1 SD. Statistical significance was assessed using two-way ANOVA of log-transformed data, followed by FDR correction for multiple comparisons.

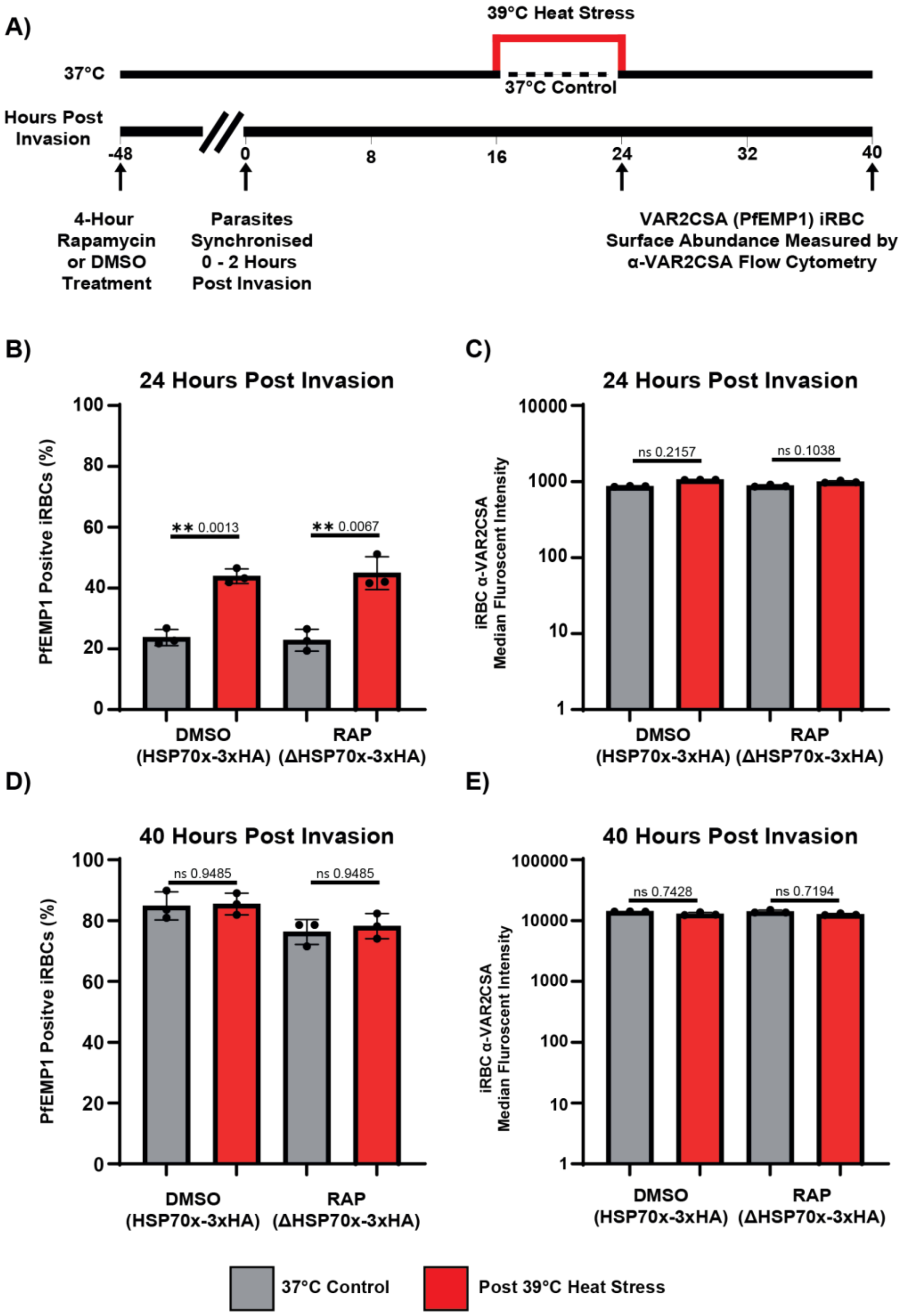


**Figure 3 - Supplement 8. The exported protein HSP70x is not required for the trafficking of PfEMP1 onto the infected cell's surface under normal or febrile temperatures.** A) Parasites were either DMSO or rapamycin (RAP) treated (four hours) to conditionally knockout HSP70x (PF3D7_0831700), parasites were later synchronised to a two-hour invasion window. Parasites were heat stressed at 39 °C between 16-24 hpi or kept at 37 °C for the full duration. At 24 and 40 hpi the presence of VAR2CSA was measured on the iRBCs surface by flow cytometry. B) At 24 hpi there was no change in the percentage of PfEMP1 positive iRBCs in the RAP compared to DMSO. There was a significant increase in the number of infected red blood cells positive for VAR2CSA following heat stress regardless of DMSO or RAP treatment. D) At 40 hpi there was no difference in percentage VAR2CSA positive iRBCs regardless of RAP or heat treatment. The α-VAR2CSA median fluorescent intensity (MFI) was not significantly different with heat or rapamycin treatment but does increase 10-fold from 24 hpi (C) to 40 hpi (E). N=3 biological replicates. Error bars displayed are ± 1 SD. Statistical significance was determined using two-way ANOVA on log-transformed data with FDR correction.

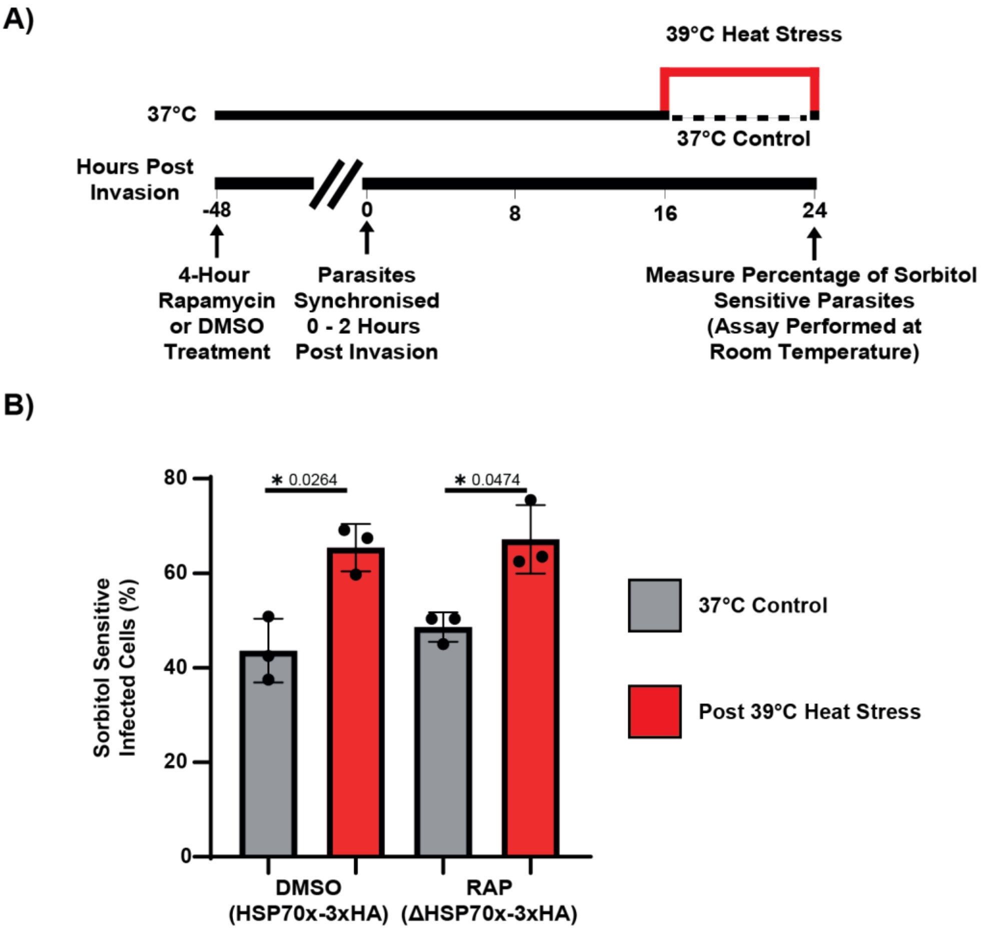


**Figure 3 - Supplement 9. HSP70x is not required for functional PSAC activity at 24 hpi under normal or elevated temperatures.** A) Parasites were either DMSO or rapamycin (RAP) treated (four hours) to conditionally knockout HSP70x (PF3D7_0831700), parasites were later synchronised to a two-hour invasion window. Parasites were heat stressed at 39 °C between 16-24 hpi or kept at 37 °C for the full duration. At 24 hpi aliquots of cells were treated with PBS or Sorbitol before parasitaemia was measured by flow cytometry. B) The reduction in parasitaemia with sorbitol treatment (Sorbitol/PBS) was measured and showed that in the absence of HSP70x (RAP) there was no change in percentage of sorbitol sensitive cells at 37 °C. Furthermore, there was an increase in the percentage of sorbitol sensitive following heat stress. Data represent n = 3 biological replicates. Error bars indicate ±1 SD. Statistical significance was determined using two-way ANOVA on log-transformed data with FDR correction.

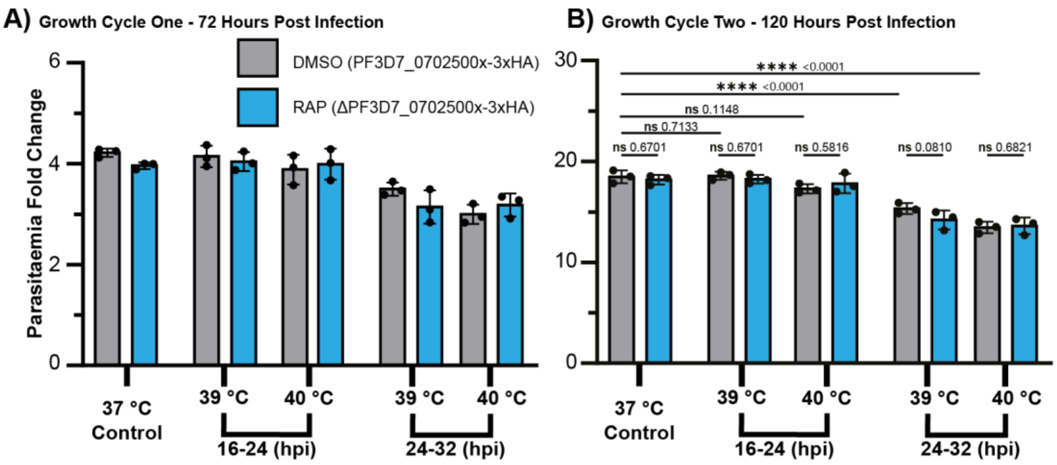


**Figure 3 - Supplement 10. PF3D7_0702500 conditional knockout parasites have normal growth at 37 °C and elevated temperatures.** Parasites were either DMSO or rapamycin (RAP) treated (four hours) to conditionally knockout PF3D7_0702500, parasites were later synchronised to a two-hour invasion window. Parasites were heat stressed at 39 °C or 40 °C between 16-24, 24-32 hpi or kept at 37 °C for the full duration. 40 °C heat stress between 24-32 hpi significantly reduced growth as shown in the fold change in parasitaemia in the first (A) and second cycle (B) following heat stress. However, the absence of PF3D7_0702500 (RAP treated) did not have a significant impact on the growth of the parasites following any of the heat stress conditions tested compared to the wildtype resembling (DMSO treated) strain. Data represent N=3 biological replicates, with error bars showing ±1 SD. Statistical significance was assessed using two-way ANOVA of log-transformed data, followed by FDR correction for multiple comparisons.

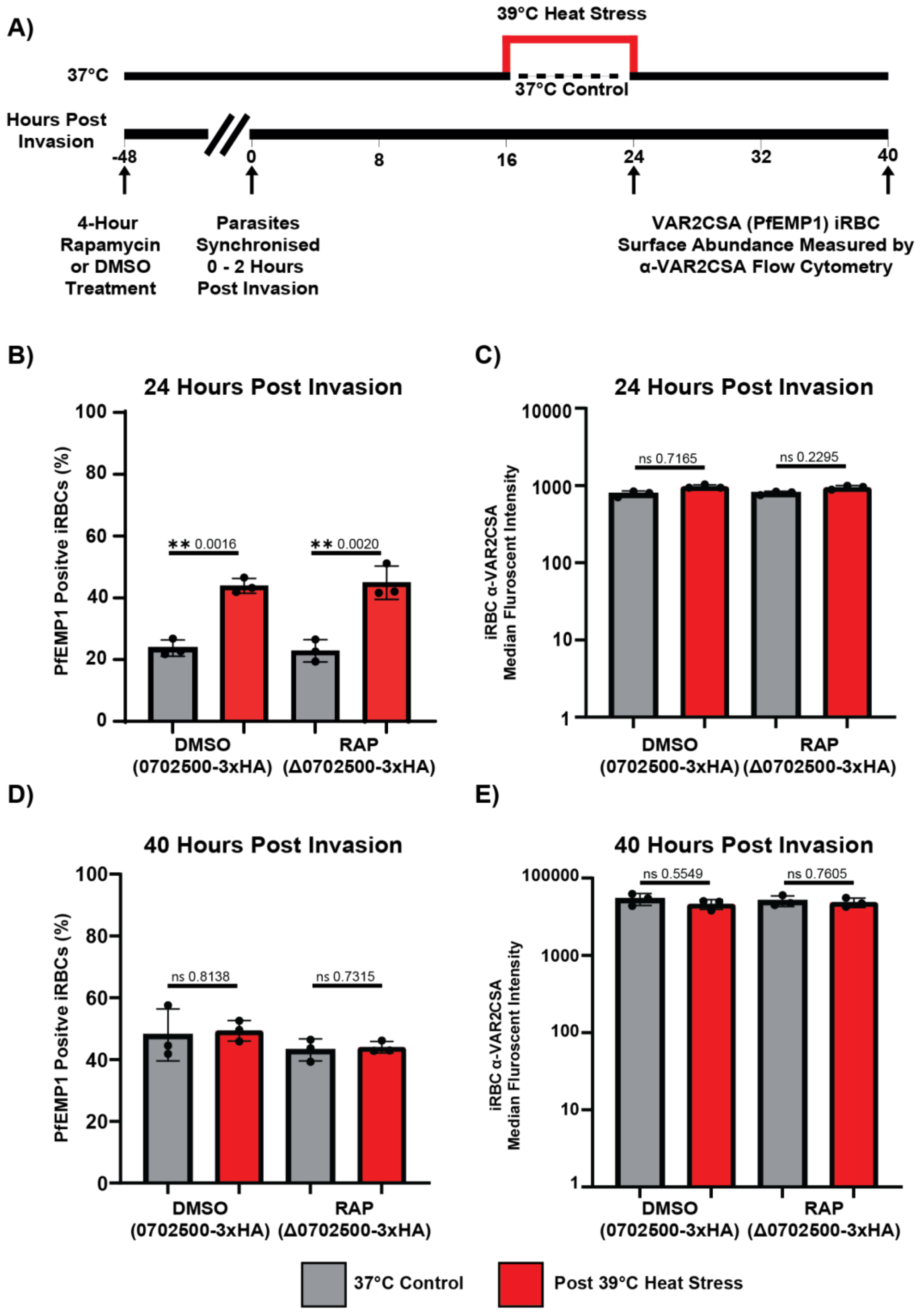


**Figure 3 - Supplement 11. The exported protein PF3D7_0702500 is not required for the trafficking of PfEMP1 onto the infected cell's surface under normal or febrile temperatures.** A) Parasites were treated with either DMSO or rapamycin (RAP) for four hours to conditionally knock out PF3D7_0702500. Parasites were subsequently synchronised to a two-hour invasion window. Heat stress was applied at 39 °C from 16–24 hours post-invasion (hpi), while control parasites were maintained at 37 °C. The presence of VAR2CSA on the iRBC surface was assessed by flow cytometry at 24 and 40 hpi. (B) At 24 hpi, there was no change in the percentage of PfEMP1-positive iRBCs in RAP-treated parasites compared to DMSO. Heat stress significantly increased the proportion of VAR2CSA-positive iRBCs, irrespective of treatment. (D) At 40 hpi, there was no difference in the percentage of VAR2CSA-positive iRBCs with either RAP or heat treatment. (C, E) Median fluorescent intensity (MFI) of anti-VAR2CSA staining was not significantly affected by heat or RAP treatment. Data represent n = 3 biological replicates. Error bars indicate ±1 SD. Statistical analysis was performed using two-way ANOVA with FDR correction. Data were log-transformed prior to analysis.

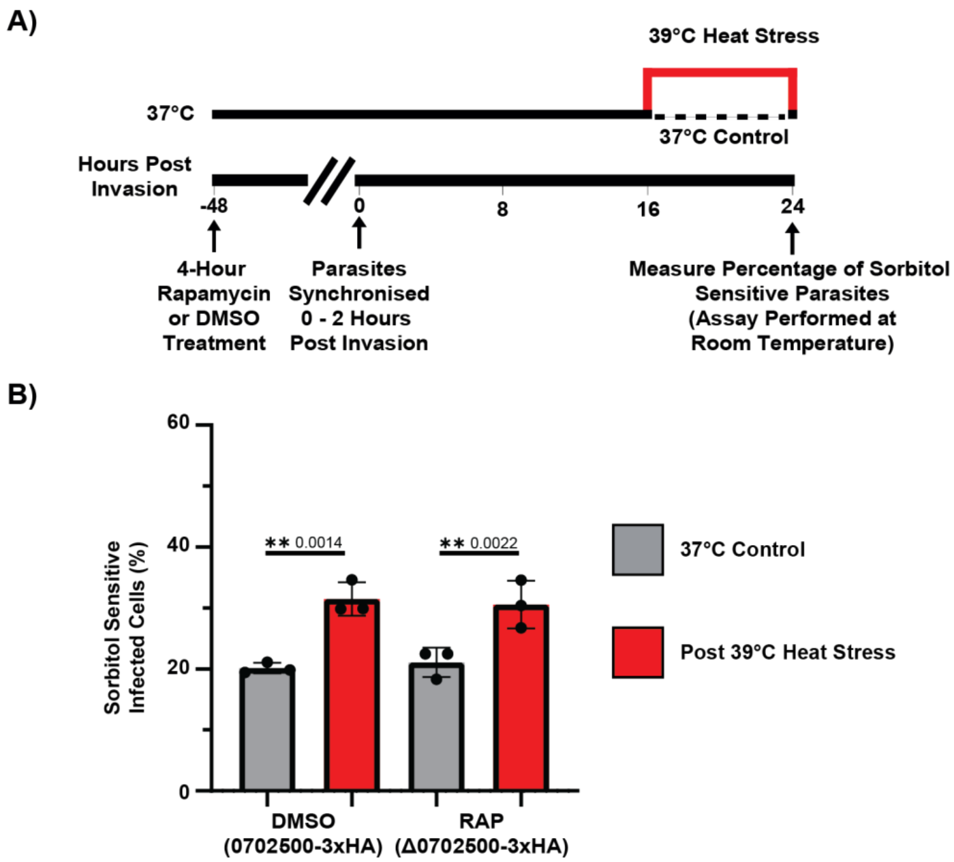


**Figure 3 - Supplement 12. PF3D7_0702500 is not required for functional PSAC activity at 24 hpi under normal or elevated temperatures.** A) Parasites were treated with either DMSO or rapamycin (RAP) for four hours to conditionally knock out PF3D7_0702500. Parasites were subsequently synchronised to a two-hour invasion window. Heat stress was applied at 39 °C between 16–24 hours post-invasion (hpi), or parasites were maintained at 37 °C. At 24 hpi, aliquots of cells were treated with either PBS or sorbitol, and parasitaemia was assessed by flow cytometry. (B) The reduction in parasitaemia following sorbitol treatment (Sorbitol/PBS) was a surrogate for functional PSAC activity. At 37 °C, deletion of PF3D7_0702500 (RAP treatment) had no effect on the percentage of sorbitol-sensitive cells. Heat stress led to an increased proportion of sorbitol-sensitive infected red blood cells in both the absence and presence of PF3D7_0702500. Data represent n = 3 biological replicates. Error bars indicate ±1 SD. Statistical significance was determined using two-way ANOVA on log-transformed data with FDR correction.

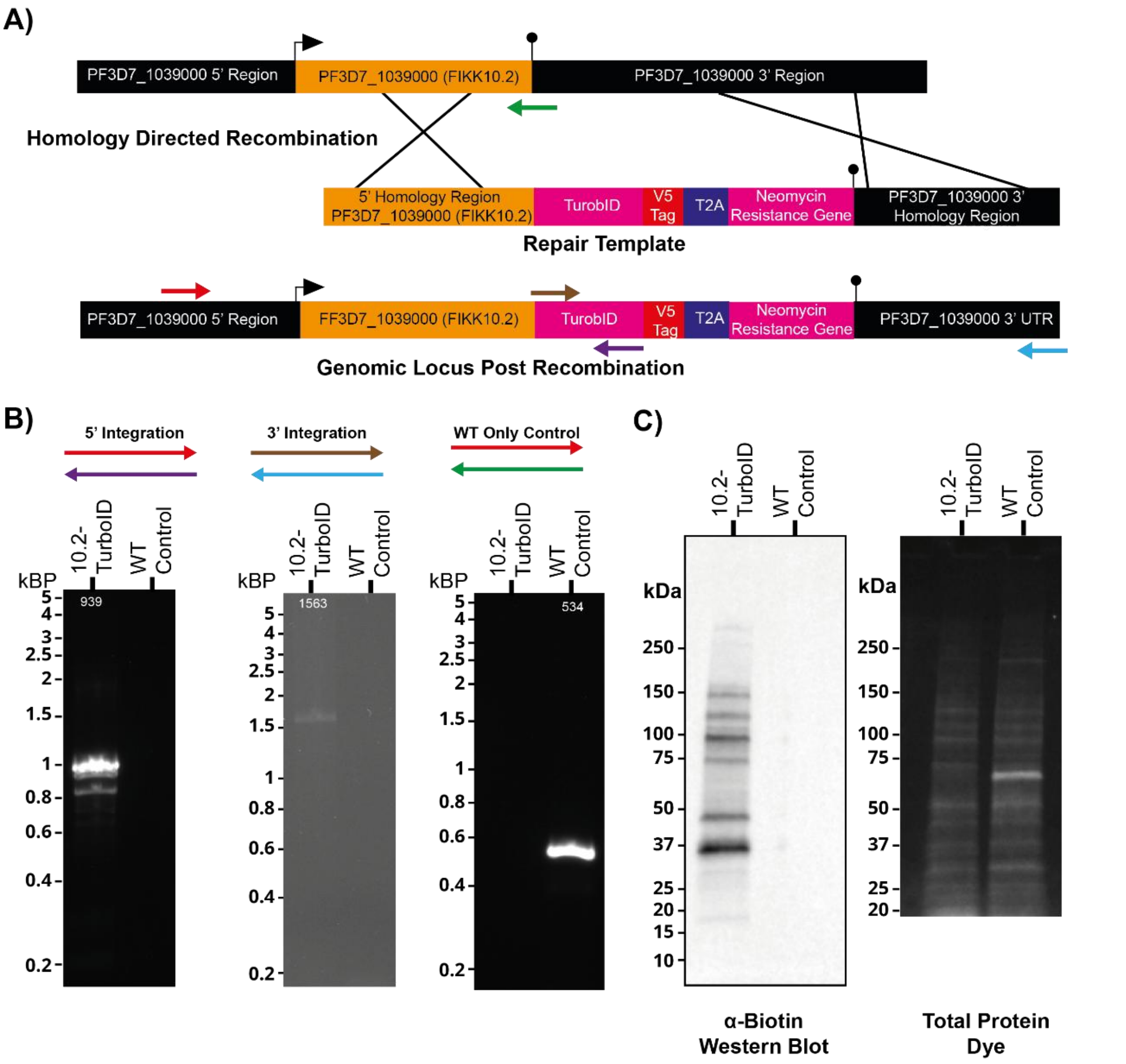


**Figure 4 – Supplement 1. Generation and validation of FIKK10.2-TurboID *P. falciparum* transgenic strain.** A) An FIKK10.2-TurboID endogenous protein fusion was generated by CRISPR/Cas9 excision and homologous repair. Primer binding sites indicated by the red and blue arrows are outside the 5' and 3' repair template homology regions. (B) 5' integration, 3' integration and the absence of the original (WT) parental strain was confirmed by PCR. Predicted amplicon sizes are indicated in white. DNA size marker values (kbp) are indicated to the left of the gel. C) Western blotting (α-Biotin) of percoll enriched mature stages transgenic FIKK10.2-TurboID parasites and a WT control (parental strain) grown in the presence of biotin show promiscuous biotinylation only in FIKK10.2-TurboID strain. A total protein dye was included as a comparative loading control.

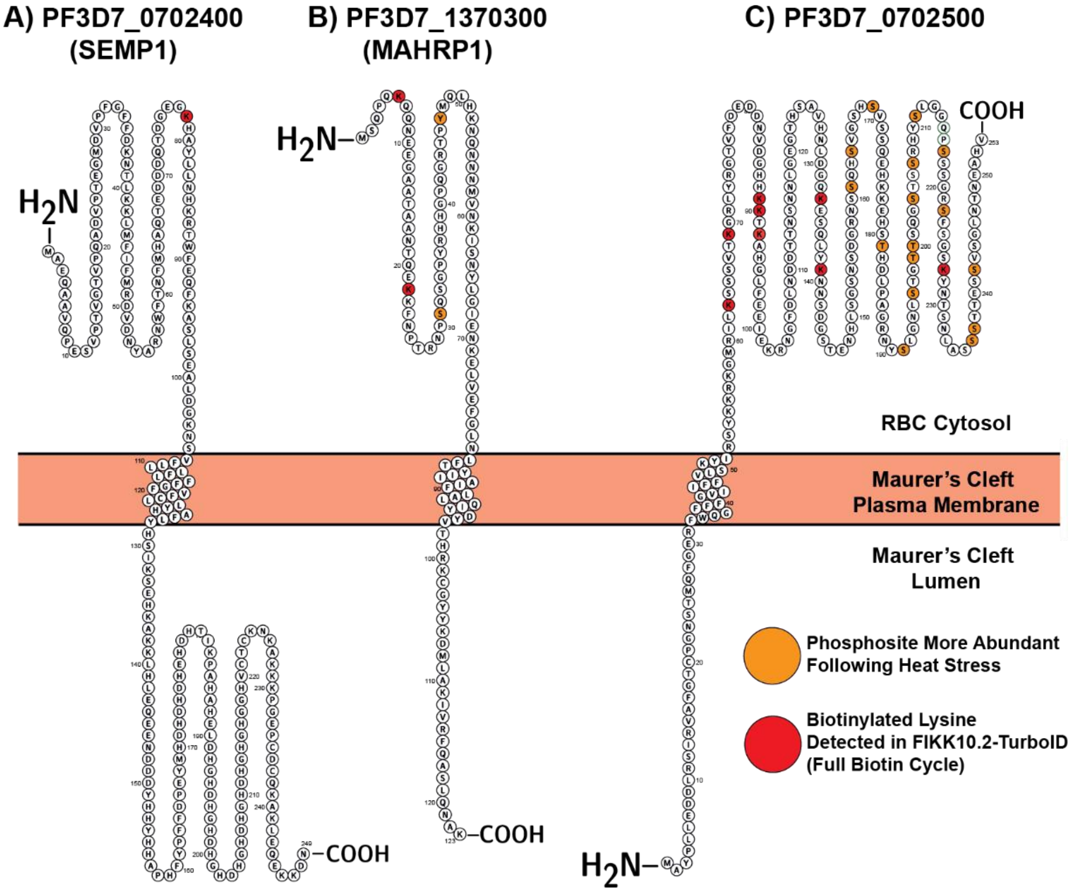


**Figure 4 – Supplement 2. Predicted Maurer's cleft protein topology from enriched phosphopeptides and biotinylated peptides.** Single letter amino acid layouts of the Maurer's cleft residing proteins (A) PF3D7_0702400 (SEMP1), (B) PF3D7_1370300 (MAHRP1) and (C) PF3D7_0702500. Predicted transmembrane domains (TMDs) are shown within the Maurer's cleft plasma membrane. N and C-termini are shown as $H_2N$ and COOH respectively. Biotinylated lysines detected within all three replicates of the FIKK10.2-TurboID (full asexual cycle condition) are highlighted in red. Phosphorylated amino acids from phosphopeptides that are more abundant following heat stress are shown in orange. All three proteins are exported into the host cell without a PEXEL motif (PNEPs) and are predicted to contain only one TMD, however, only PF3D7_0702500 is predicted to have the N-terminus within the Maurer's Cleft lumen. Single amino acid schematic was generated with the Protter Tool [86].

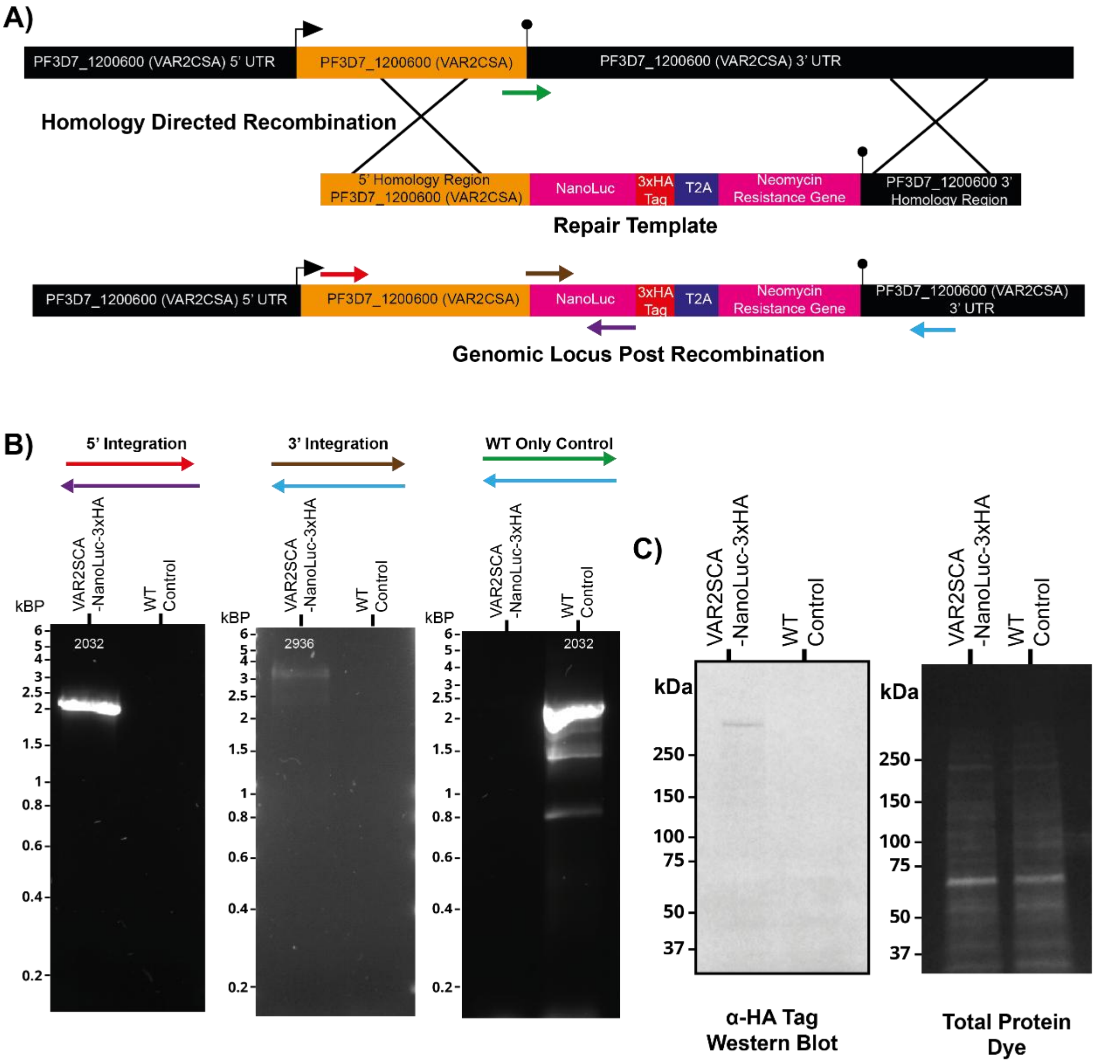


**Figure 5 – Supplement 1. Generation and validation of VAR2CSA-NanoLuc-3xHA *P. falciparum* transgenic strain.** A) VAR2CSA-NanoLuc-3xHA protein fusion was generated by CRISPR/Cas9 excision and homologous repair. Primer binding sites indicated by the red and blue arrows are outside the 5' and 3' repair template homology regions. Predicted amplicon sizes are indicated in white. (B) 5' integration, 3' integration and the absence of the original (WT) parental strain was confirmed by PCR. (C) Western blotting (α-HA Tag) of percoll enriched mature stages transgenic VAR2CSA-NanoLuc-3xHA parasites and a WT control (parental strain). A total protein dye was included as a comparative loading control. VAR2CSA-NanoLuc-3xHA has a predicted molecular weight of 379.8 kDa.

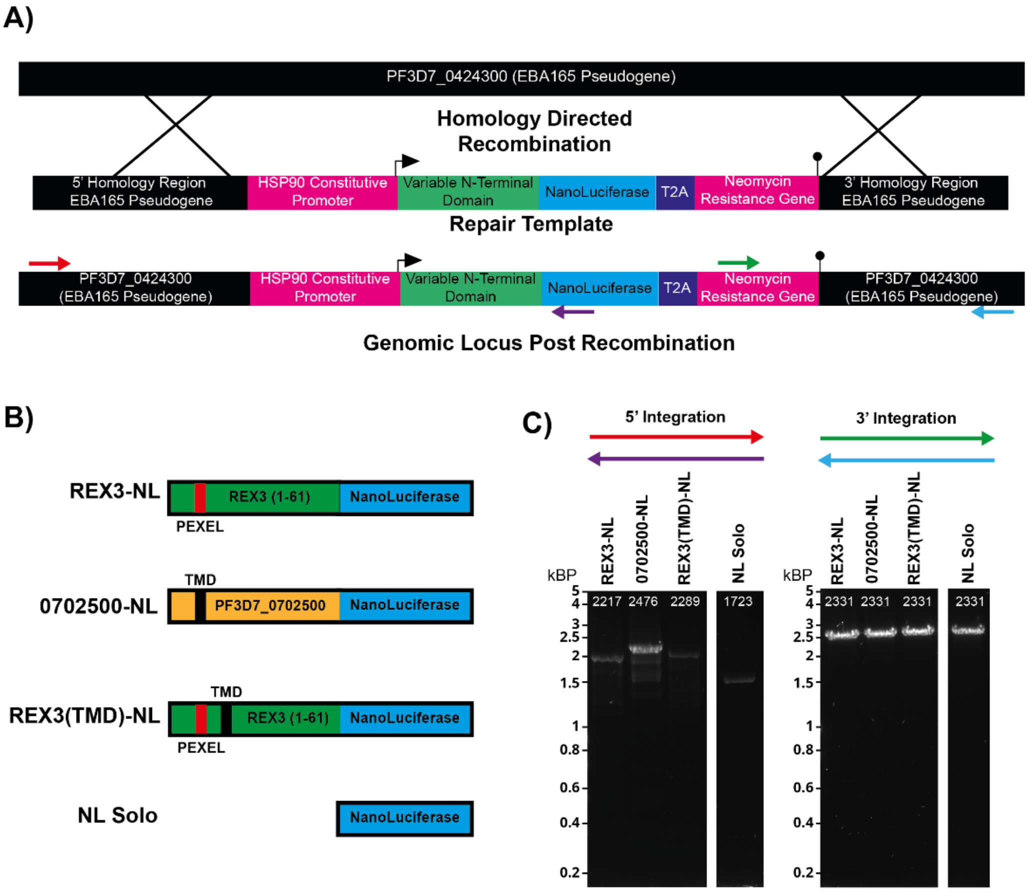


**Figure 6 – Supplement 1. Generation and validation of constitutively expressed NanoLuc reporter strains.** A) Following a CRISPR/Cas9 directed double-stranded DNA break, reporter constructs were inserted into the EBA165 Pseudogene Locus (PF3D7_0424300). Reporter constructs consisted of a constitutive promoter (HSP90, PF3D7_0708400), a variable N-terminal domain, NanoLuc and a T2A skip peptide followed by a neomycin resistance cassette. Primer binding sites indicated by the red and blue arrows are outside the 5' and 3' repair template homology regions. (B) Four distinct N-terminal variants were generated. REX3-NanoLuc (NL) comprises the first 61 amino acids of REX3, which are sufficient to mediate export [59,60]. PF3D7_0702500-NL represents a fusion of the PF3D7_0702500 N-terminus with NL. REX3(TMD)-NL refers to REX3-NL incorporating the transmembrane domain (TMD) of PF3D7_0702500. Finally, NL alone, lacking any N-terminal domain, was included. (C) Successful 5′ and 3′ integration of all four constructs into the EBA165 locus was confirmed by PCR. Predicted amplicon sizes are indicated in white.
